# Improved discovery of RNA-binding protein binding sites in eCLIP data using DEWSeq

**DOI:** 10.1101/2022.11.15.516416

**Authors:** Thomas Schwarzl, Sudeep Sahadevan, Benjamin Lang, Milad Miladi, Rolf Backofen, Wolfgang Huber, Matthias W Hentze, Gian G Tartaglia

## Abstract

Enhanced crosslinking and immunoprecipitation (eCLIP) sequencing is a powerful method for transcriptome-wide detection of binding sites of RNA-binding proteins (RBPs). However, identified crosslink sites can profoundly deviate from experimentally established functional elements of even well-studied RBPs. Current peak-calling strategies result in low replication and high false-positive rates. Here, we present the R/Bioconductor package *DEWSeq* that makes full use of replicate information and size-matched input controls. We benchmarked *DEWSeq* on 107 RBPs for which both eCLIP data and RNA sequence motifs are available and were able to more than double the number of motif-containing binding regions relative to standard eCLIP processing (2.3-fold median). The improvement not only relates to the number of binding sites (e.g., 3.1-fold of known motifs for RBFOX2), but also their subcellular localisation (e.g., 1.9-fold of mitochondrial genes for FASTKD2) and structural targets (e.g., 2.2-fold increase of stem-loop regions for SLBP). DEWSeq therefore shows promise as an improved processing method for eCLIP protein–RNA interaction data.

## Introduction

RNA-binding proteins play major roles in biological processes such as splicing^1^, polyadenylation, nuclear export, subcellular localisation, transcript stabilisation and degradation as well as translation^2^. In recent years, thousands of mammalian proteins have been found to bind to RNA^3,4^, many of which have unknown RNA targets. To identify RNA sites bound by an RBP of interest, several related crosslinking and immunoprecipitation (CLIP) high-throughput sequencing methods have been developed^5^. These methods exploit the phenomenon that UV light induces covalent RNA-protein crosslinks between RNA nucleotides and protein amino acids in immediate contact with each other^6^. Over the years, several variants of CLIP-based sequencing methods have been developed: HITS-CLIP^7^ directly sequences the crosslinked RNA fragment, PAR-CLIP detects mutations induced at the crosslink site^8^, the related methods iCLIP^9^ and eCLIP^10^, as well as further derivatives such as irCLIP^11^, seCLIP^12^ and easyCLIP^13^ optimise the protocols for reverse transcription truncations for precise identification of RNA-protein crosslink sites.

Enhanced CLIP (eCLIP) introduced changes to the sequence library generation as well as a size-matched input (SMI) control to address background noise and false positives in CLIP data^10^. The ENCODE Consortium has used eCLIP to generate the largest coherent public set of CLIP data, covering 150 RNA-binding proteins in two cell types (HepG2 and K562), processed with the computational peak-calling analysis pipeline CLIPper^10^. Detected binding sites were compared against SMI controls individually for each replicate and extended by 50 nucleotides from their 5’ end for functional analyses, with the reasoning that the 5’ end of a peak represents the crosslink site^14^. An analysis of these data shows the relatively low reproducibility of reported binding sites between replicates (Fig. 1a). While RBPs such as the SBDS Ribosome Maturation Factor (SBDS), NOP2/Sun RNA Methyltransferase 2 (NSUN2), and Small RNA Binding Exonuclease Protection Factor La (SSB) show almost perfect reproduction of the respective binding sites, Transforming Growth Factor Beta Regulator 4 (TBRG4), Splicing Factor 3b Subunit 1 (SF3B1), and WD Repeat Domain 3 (WDR3) binding sites display high replicate to replicate variation. More recently, a ‘*CLIPper reproducible*’ (*CLIPper*_*Rep*._) dataset was introduced, featuring only binding sites that were identical at the base level in both replicates^14^. This approach greatly reduced the number of reported binding sites. However, it raises the question whether better data analysis approaches exist. A particular challenge in the analysis of eCLIP data is that the measured crosslink peaks can be at an offset from the RBP’s actual binding regions, which can result from an RBP’s particular structure and physicochemical crosslinking behaviour. CLIP methods are often tested against classical RBPs such as the RNA Binding Fox-1 Homolog 2 (RBFOX2) or splicing factor Heterogeneous Nuclear Ribonucleoprotein C (hnRNPC) with well-known binding sites and RNA sequence motifs^10,13,15^. RBFOX2’s crosslink site pattern around discovered binding sites is strongest at its known RNA sequence motifs (Fig. 1b). Similarly, hnRNPC contacts RNA in a motif- and position-dependent context, displaying bell shaped crosslink distribution around the expected binding site^16^. Both cases support the use of traditional peak-callers. However, other RBPs display profound divergences between their biological binding sites and their crosslink behaviours in terms of positioning and shape of the truncation/crosslink sites (Fig. 2 and Supplementary Fig. 1a,b): For example, the stem-loop binding protein (SLBP) protein, a protein which binds conserved 3’UTR stem-loop structures in histone genes^17^, shows systematic crosslink site enrichment upstream of the actual stem-loop (Fig. 1c). CSTF2, known to interact with the AAUAAA polyadenylation signal^18^, conversely displays crosslink enrichment downstream of its binding motif (Fig. 1d), while U2AF2 binds either directly at or downstream of its uridine/cytidine-rich motifs^19^ (Fig. 1e). Other RBPs such as HNRNPL^20^ (Fig. 1f), CPEB4^21^ (Fig. 2 and Supplementary Fig. 1a,b) or the non-classical RBP ENO1 show different crosslinking behaviour^10,22^. HNRNPL crosslink sites are in fact depleted at its known sequence motif (Fig. 1f and Supplementary Fig. 1a,b). Generally, the crosslink sites are enriched at or in proximity of the binding motif (Fig. 2 and Supplementary Fig. 1a,b). The shift of crosslink site peaks was also evident in some iCLIP data: for eIF4A3, an exon junction complex subunit with well-known binding site locations, the bell-shaped crosslink site curve was shifted by >10 nt compared to other exon junction complex proteins^23,24^. Without prior knowledge of an RBP’s behaviour, such shifts in positioning and varying crosslink site behaviour are likely to lead to misinterpretation of the binding sites.

**Figure 1.**
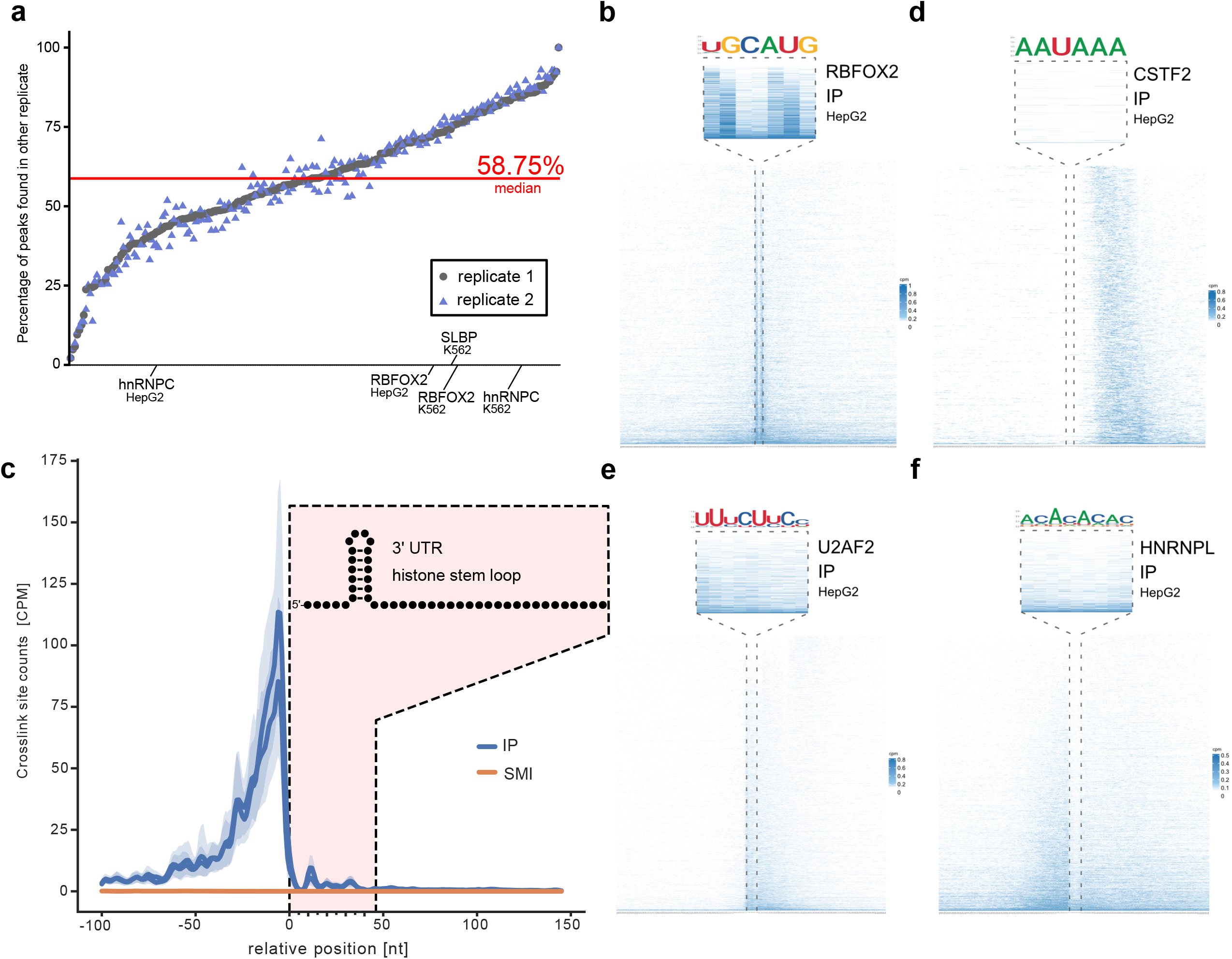
eCLIP crosslink sites around functional elements. (**a**) Reproducibility of binding sites between ENCODE eCLIP data sets replicate 1 and 2. A binding site is counted as reproducible if at least 1 nucleotide overlaps with a binding site called in the other replicate. (**b**) Example of RBFOX2 crosslink site distribution for ENCODE data set ENCSR756CKJ (K562) and ENCSR987FTF (HepG2) relative to known RBFOX2 UGCAUG motif. (**c**) Crosslink site distribution of SLBP eCLIP data set on 34 histone genes relative to known 3’ UTR histone stem-loop (ENCODE eCLIP data set ENCSR483NOP, K562 cell line). (**d**) CSTF2 crosslink site distribution for ENCODE eCLIP data set ENCSR384MWO (HepG2 cell line) relative to known AAUAAA polyadenylation signal. (**e**) U2AF2 crosslink site distribution for ENCODE eCLIP data set ENCSR202BFN (HepG2 cell line) relative to uridine/cytidine-rich motifs. (**f**) HNRNPL crosslink site distribution for ENCODE eCLIP data set ENCSR724RDN (HepG2 cell line) relative to CA repeat motifs.

**Figure 2.**
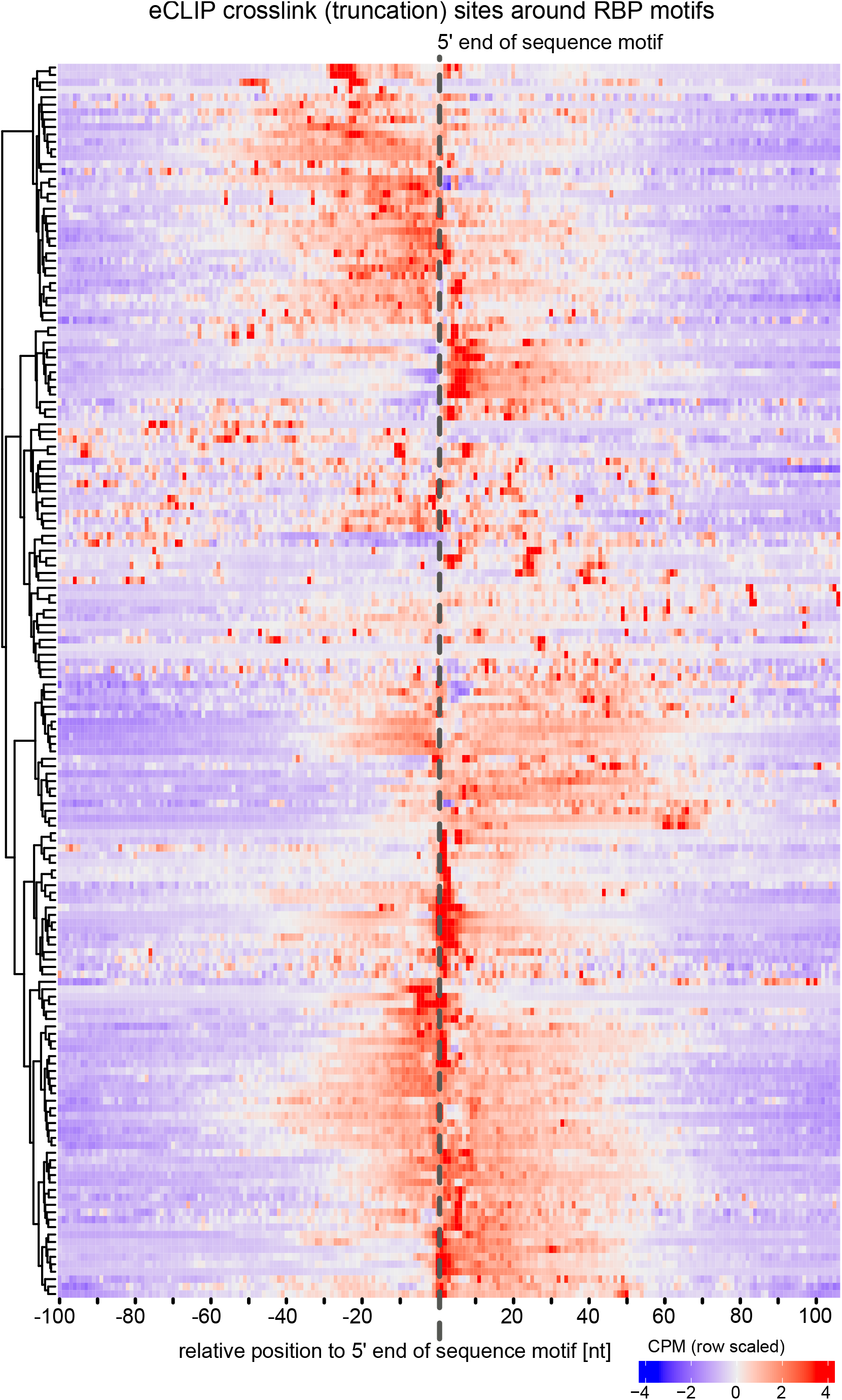
eCLIP crosslink sites around known motifs. Heatmap of eCLIP crosslink (truncation) sites around experimentally derived RNA sequence motifs for 107 RBP from ENCODE. Each row displays a motif for an RNA-binding protein per eCLIP data set.

Given the varying crosslinking behaviour of each protein and the relatively low reproducibility of binding site detections in eCLIP experiments, we developed a method, *DEWSeq*, that allows accurate and robust identification of RBP interactions in eCLIP data by detecting regions enriched in crosslink sites compared to the control. *DEWSeq* is a sliding-window-based approach that uses single-nucleotide precision information across multiple replicates and control experiments for significance testing. To test *DEWSeq* and to facilitate a comprehensive analysis of RBP-RNA interactions, we benchmarked it on the eCLIP data for RBPs provided by ENCODE. Notably, 107 out of 150 RBPs in the dataset have known experimentally determined RNA sequence motifs, and one, SLBP, is known to recognise a specific secondary structure (histone mRNA stem-loops). We used validated RNA motifs as a proxy for the biological relevance of a given binding site. This compilation represents, to the best of our knowledge, the most comprehensive eCLIP benchmark based on known sequence motifs to date. We show that RNA binding regions identified by *DEWSeq* show a consistent improvement in sensitivity as well as specificity relative to HITS-CLIP, iCLIP and PAR-CLIP experiments.

## Results

We developed *DEWSeq* as a new R/Bioconductor statistical analysis package for the robust detection of RBP binding regions from i/eCLIP data sets. The *DEWSeq* workflow starts from the output of an accompanying Python package for post-processing i/eCLIP alignment files, *htseq-clip*^*25*^, which extracts crosslink site counts at single-nucleotide positions adjacent to the end of reads, flattens annotation of multiple transcripts and uses sliding windows to count and aggregate crosslink sites. *DEWSeq* performs one-tailed significance testing using *DESeq2*^*26*^, result summarisation and binding site visualisation (Supplementary Fig. 2).

Similar to the *csaw* package^27^ for ChIP-seq data, *DEWSeq* incorporates biological variation with significance testing, which reduces the false discovery rate^27^, with the difference that *DEWSeq* is tailored to single-nucleotide position data. *DEWSeq* successfully utalises SMI as input control, while previously it was only used in a rank-based metric for determining the specificity of binding^28^. 223 ENCODE eCLIP datasets covering 150 RBPs cell types HepG2 and K562 were processed and analysed with *htseq-clip/DEWSeq*.

### Sequence motif-based evaluation strategy

We compared DEWSeq’s results to the *CLIPper* method used by the ENCODE Project, which uses peak-calling on two individual replicates compared against a single SMI control. We extended each peak 50 nt in the 5’ direction, as first introduced by the authors specifically for motif-based analyses^14^, which is referred to as *CLIPper original (CLIPper*_*Orig*._) in our study. This dataset was improved on by the authors to produce *CLIPper*_*Rep*._, which is the subset of *CLIPper*_*Orig*._ peaks that are reproducible at the nucleotide level across both replicates^10,14^. An overview of the RNA sequence motif-based benchmarking strategy we adopted in our study is shown in Fig. 3a.

**Figure 3.**
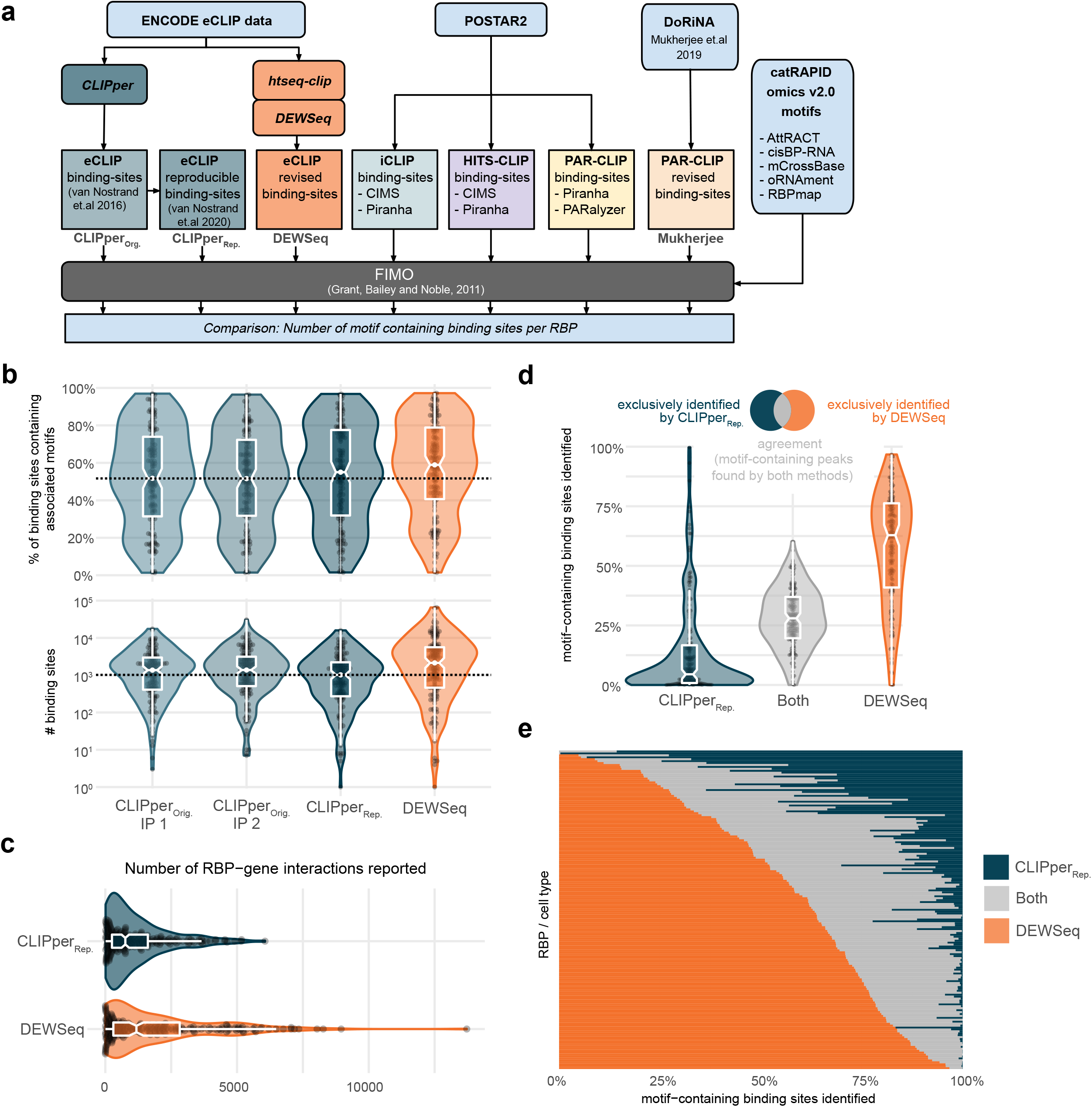
Overview and results for benchmarking workflow on ENCODE eCLIP data sets for proteins with known RNA sequence motifs. (**a**) ENCODE eCLIP data sets were reanalysed with *DEWSeq* and compared to ‘*CLIPper original*’ (*CLIPper*_*Orig*.._)^*10,29*^ and ‘*CLIPper reproducible*’ (*CLIPper*_*Rep*._) data set^14^ analysis. Other CLIP binding sites from iCLIP, HITS-CLIP and PAR-CLIP were extracted from POSTAR2. Additional PAR-CLIP data sets from DoRiNA^32^ were included in the analysis. Binding sites from all data-sets were analysed with FIMO^30^ using known RNA sequence motifs from catRAPID omics v2.0^29^. (**b**) Top panel shows violin and boxplots of the number of motif-containing binding sites in datasets detected with FIMO and catRAPID omics v2.0 motifs. Bottom panel violin and boxplots show the percentage of motif-containing binding sites to the total number of binding sites for each method. (**c**) Number of reported RBP-gene interactions. (**d**) Exclusiveness of motif-containing binding sites for data sets with known motifs. Left shows binding sites exclusive for *CLIPper*_*Rep*._ data set, the middle the binding sites which were found in both, and right for sites found exclusively in *DEWSeq*. (**e**) Heatmap of exclusiveness and overlap for motif-containing *CLIPper*_*Rep*._ and *DEWSeq* binding sites.

As a proxy for the likely biological relevance of the identified binding sites, we obtained known experimentally determined RNA sequence motifs of 6 nucleotides or longer from *cat*RAPID omics v2.0^29^ (Supplementary File 1). This curated motif dataset covered 107 of the 150 RBPs for which ENCODE eCLIP data were available.

To identify positions of known motifs within binding regions identified by *CLIPper* and *DEWSeq* from ENCODE datasets, we used FIMO from the MEME suite of motif analysis software^30^. To compare these results to orthogonal datasets beyond eCLIP, we obtained iCLIP, HITS-CLIP, PAR-CLIP binding sites determined using different peak callers in the POSTAR2 database^31^ as well as a separate dataset of author-processed PAR-CLIP binding sites^32^, and scanned for motifs using FIMO. For each method, we then estimated the accuracy of binding site detection by calculating the proportion of reported binding sites that contained at least one expected sequence motif for the RBP of interest.

### Performance comparison between DEWSeq and CLIPper

Slightly over half (51.8%) of the *CLIPper*_*Orig*._ binding sites contained a known sequence motif for the RBP under investigation (median across RBPs, cell types and replicates, Fig. 3b bottom). *CLIPper*_*Rep*._ slightly increased the motif-containing binding site fraction to 55.0%, but at the expense of reducing the total number of motif-containing regions identified from 1,366 to 1,021 median binding sites per RBP and cell type (Fig. 3b top). Conversely, while *DEWSeq* binding sites showed a further increased motif-containing rate (58.9%), *DEWSeq* also notably identified a total number of motif-containing binding sites (median 2,137) that was markedly higher than both *CLIPper*_*Rep*._ and the less stringent *CLIPper*_*Orig*._ set (a 2.25-fold and 1.81-fold median improvement, respectively). Complete results from this analysis are provided in Supplementary Table 1. Thus, *<u>DEWSeq</u>*<u>achieves an increase in the number of detected binding sites without reducing the proportion of motif-containing sites,</u> nor with any apparent systemic bias towards gene regions or gene types (Supplementary Fig. 3). We thus posit that the detection quality is on median at least as good as that of the *CLIPper*_*Rep*._ approach, while the detection rate is more than twice as high. At the gene level, *DEWSeq* consequently increases the discovery of RBP-RNA interactions from a median of 760 to 1,181 genes per RBP and cell type (Fig. 3c).

### Motif exclusiveness analysis

Following the motif-containing binding site analysis, we assessed what proportion of binding sites that *DEWSeq* and *CLIPper* have in common and also the proportion of binding sites that are exclusively found by one of the respective methods. We compared *CLIPper*_*Rep*._ and *DEWSeq* as the best-performing representatives for the two analysis strategies (data for all runs are provided in Supplementary Table 2, Sheet 1), and focussed on the binding sites with known motifs. In this analysis, all detected motifs were classified based on whether they were detected by both methods (labelled ‘Both’), or exclusively by only one of the methods (indicated either as *CLIPper*_*Rep*._ or *DEWSeq*, respectively). For each RBP, the percentage of motif-containing peaks in each group was calculated (Fig. 3d,e). A substantial fraction of motif-containing binding regions was detected exclusively by *DEWSeq* (median across RBP–cell type experiments: 62.9%). *DEWSeq* and*CLIPper*_*Rep*._ showed good agreement on binding sites for some RBPs such as EFTUD2 (47.4% in K562 and 37.2% in HepG2 cells), while for other RBPs such as AKAP1 the agreement was high in one cell type (K562, 34.7%), but not in the other (HepG2, 11.4%) (Fig. 3d). Overall, the median fraction of motif-containing binding regions exclusively detected by *DEWSeq* (62.9%) greatly exceeded the fraction agreed on by both methods (28.0%) and those found exclusively by *CLIPper*_*Rep*._ (4.5%) (Fig. 3d,e and Supplementary Table 2, Sheet 1).

### Results for specific RBPs with well-defined biological roles

#### RBFOX2

RBFOX2 (RNA Binding Fox-1 Homolog 2) is an alternative splicing regulator that binds to UGCAUG motifs^33^. It is regularly used for benchmarking in CLIP manuscripts^10^. Here, we compared the number of RBFOX2 binding regions containing the UGCAUG motif reported by *CLIPper*_*Rep*._, *DEWSeq* or both. Fig. 4a shows the total number of regions reported and the number of regions including the UGCAUG motif. The fraction of motif-containing regions per cell line (HepG2 and K562) are similar for both *CLIPper*_*Rep*._ and *DEWSeq* results: 61.2% and 60.2% in HepG2 cell lines and 44.9% and 46.7% in K562 cell lines. However, a striking difference can be seen for the number of motif-containing regions reported by *DEWSeq* as compared to *CLIPper*_*Rep*._. In the HepG2 cell line, *CLIPper*_*Rep*._ reported 3,223 UGCAUG motif-containing regions compared to 8,410 motif-containing regions for *DEWSeq*, and in K562 cells *CLIPper*_*Rep*._ reported 1,204 motif-containing regions in comparison to 4,257 from *DEWSeq*, indicating a substantial improvement in sensitivity when using DEWSeq (Supplementary Table 3, Sheet 1).

**Figure 4.**
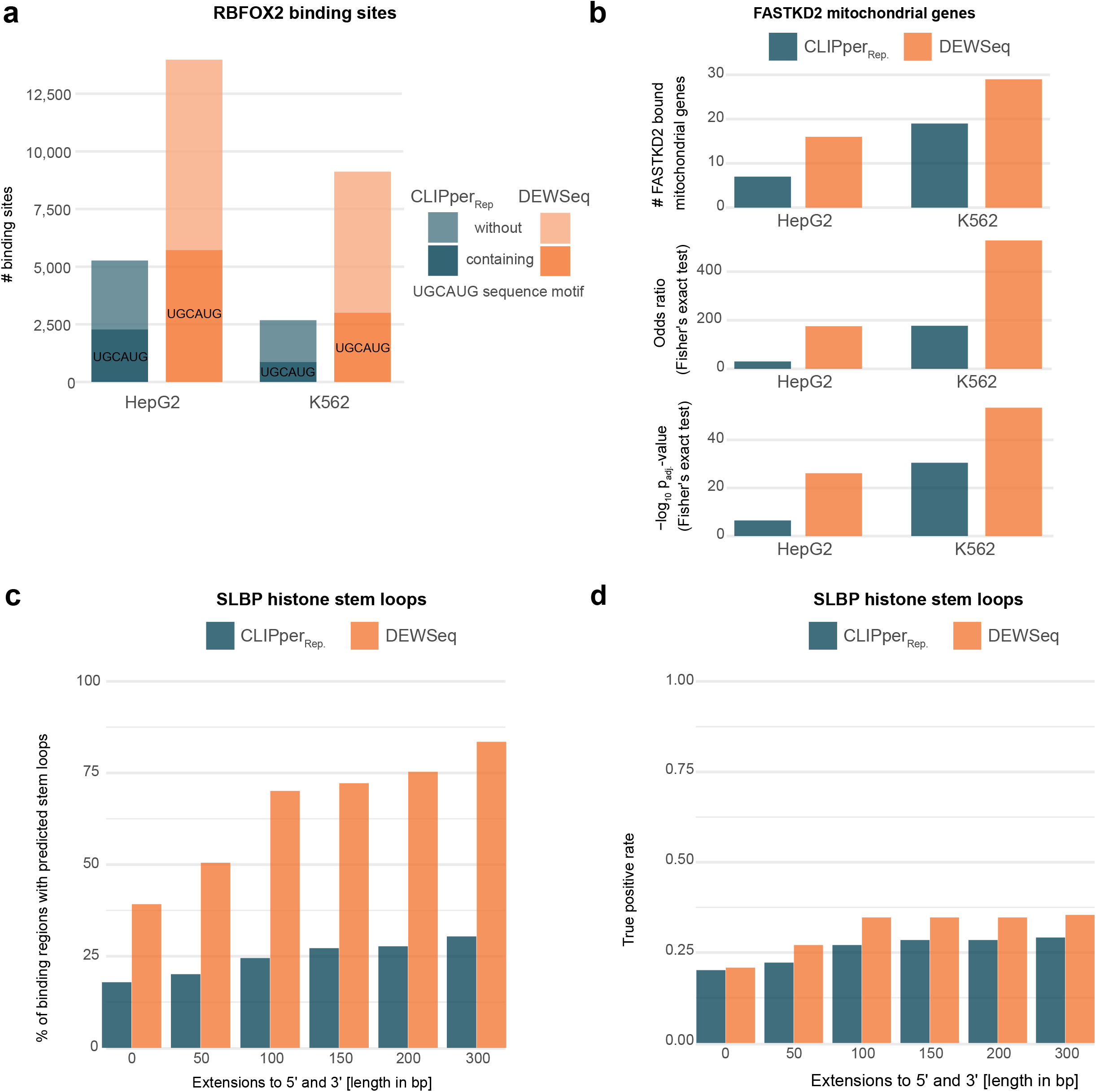
Binding site comparisons. (**a**) Comparison of RBFOX2 binding regions and RBFOX2 binding regions containing TGCAGT motif in *‘CLIPper reproducible’* (*CLIPper*_*Rep*._) and *DEWSeq*. (**b**) FASTKD2 binding site enrichment for mitochondrial genes compared to other chromosomal locations for *CLIPper*_*Rep*._ and *DEWSeq* binding sites. Supplementary Table 3, Sheet 2 contains the complete enrichment analysis results across all chromosomes for all FASTKD2 samples. (**c**) SLBP stem-loops found with 3’ and 5’ Extensions of *DEWSeq* and *CLIPper*_*Rep*._ binding sites. Left panel shows the percentage of binding regions containing the predicted stem-loops. (**d**) True positive rate (sensitivity) with respect to reference Histone 3’ UTR stem-loop regions retrieved from Rfam database (for Rfam id: RF00032).

#### FASTKD2

FASTKD2 (FAST kinase domain-containing protein 2) is a mitochondrial RBP that has been shown to interact with a defined set of mitochondrial transcripts^34,35^. In this analysis, we compared the number of FASTKD2-bound regions of *DEWSeq* against *CLIPper*_*Rep*._ results. In HepG2 cells, *CLIPper*_*Rep*._ reports 7 out of 451 bound genes as mitochondrial, compared to *DEWSeq* with 16 out of 268 bound genes being mitochondrial (Fig. 4b). A similar pattern emerges in the K562 cell line, where the numbers for *CLIPper* and *DEWSeq* were 19 out of 364 and 29 out of 426, respectively. Fisher’s Exact test confirmed a significant enrichment in the number of FASTKD2-bound mitochondrial genes reported by *DEWSeq* compared to *CLIPper*_*Rep*_ results in both cell lines (Fig. 4b, Supplementary Table 3, Sheet 2).

#### SLBP

SLBP (Stem-Loop Binding Protein) is an RBP that binds to a conserved stem-loop structure motif at the 3’ end of mRNAs that encode replication-dependent histones^2,36,37^. To the best of our knowledge, it represents the only RBP included in the ENCODE project that recognises a secondary structure motif. We scanned SLBP binding regions and surroundings (binding site were extended with 50, 100, 150, 200, and 300 nt in both 5’ and 3’ direction) from both *CLIPper*_*Rep*._ and *DEWSeq* results for the SLBP stem-loop structure binding site using Infernal suite and Histone 3’ UTR stem-loop covariance model (Rfam: RF00032).

A higher proportion of *DEWSeq* binding sites (39.2%) contain predicted SLBP binding structures, as compared to *CLIPper*_*Rep*._ binding regions (17.9%) (Fig. 4c). This trend becomes more pronounced with the extension of binding regions in both 5’ and 3’ directions (Fig. 4c and Supplementary Fig. 4), as *DEWSeq* discovers increasingly more stem-loops: <u>83.5% of detected binding sites are in proximity to histone stem-loops, whereas only 30.4% of *CLIPper*_*Rep*_. binding sites are in the vicinity of known targets</u>, suggesting a significant decrease of false positives for *DEWSeq*.

Further, we calculated the true positive rate (sensitivity) of these predicted stem-loop structures using Histone 3’ UTR stem-loop annotations from the Rfam database as a reference set (Fig. 4d). *DEWSeq* without extension shows a marginal increase in true positive rate compared to *CLIPper*_*Rep*._ (from 0.201 to 0.208), with slight improvement in extensions (Supplementary Table 4).

In addition to the increased presence of expected stem-loop structures, we also noted that *CLIPper*_*Rep*._ identified binding of SLBP to mRNAs deriving from a total of 44 histone genes (66.7% of its target genes being histones), while *DEWSeq* identified binding of SLBP to a total of 53 histone mRNAs (71.6% of its target genes being histones). For reference, the HGNC histone gene set contains a total of 118 genes.

### Evaluation of eCLIP compared to iCLIP, HITS-CLIP and PAR-CLIP

To validate newly discovered binding sites, we used the motif exclusivity benchmark to compare eCLIP binding sites assigned by *DEWSeq* and *CLIPper*_*Rep*._, respectively, to sites from iCLIP, HITS-CLIP and PAR-CLIP protocols retrieved from the POSTAR2^31^ and DoRiNA^32^ databases. To address differences in detection methods for these different CLIP protocols, we employed multiple established analysis methods: Piranha^38^ and CIMS^39^ for iCLIP and HITS-CLIP data, and Piranha^38^, PARalyzer^40^ as well as Mukherjee^32^ for PAR-CLIP data.

Fig. 5a-c left panels show the comparison of *CLIPper*_*Rep*._ results to iCLIP, HITS-CLIP and PAR-CLIP, and right panels show the comparison of *DEWSeq* results to the same. eCLIP yields more motif-containing binding sites overall, with a substantially higher number identified by *DEWSeq* compared to *CLIPper*_*Rep*._ (Fig. 5a-c, Supplementary Fig. 6a-c and Supplementary Table 2, Sheet 2). Interestingly, the overlaps of binding sites found in either eCLIP and iCLIP, HITS-CLIP, or PAR-CLIP are modest compared to binding sites detected in one of the protocols alone. *DEWSeq* recovers a median of 60 (2.1% of total) motif-containing binding sites found by other methods, compared to a median of 18 (1.0% of total) for *CLIPper*_*Rep*._ (Fig. 5d and Supplementary Fig. 6d).

**Figure 5.**
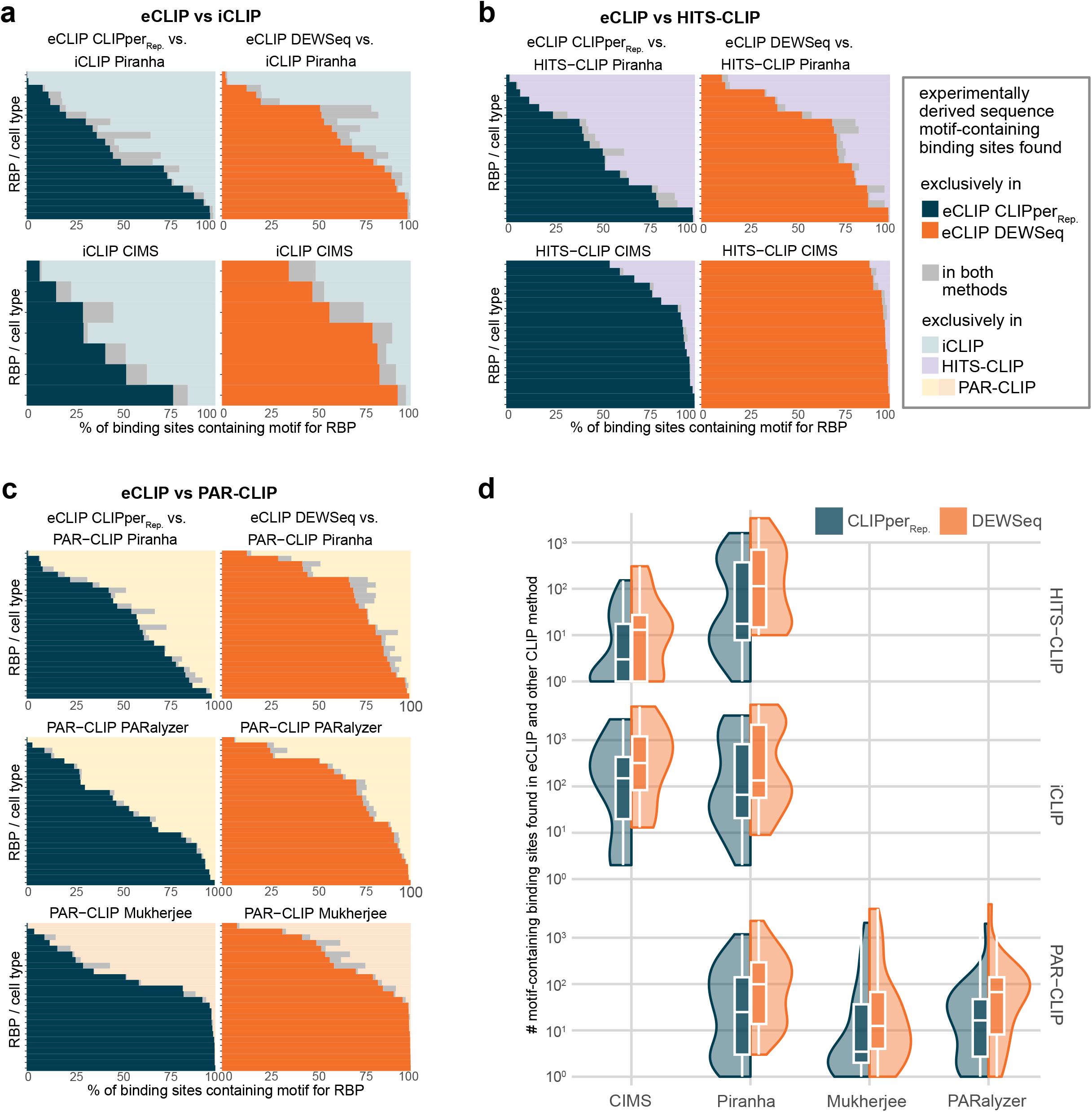
Binding site exclusiveness. Comparison of motif-containing binding sites from various CLIP datasets against eCLIP *‘CLIPper reproducible’* (*CLIPper*_*Rep*._) and *DEWSeq* results. The comparisons are for common RBPs (in ENCODE dataset and these methods) with motifs in catRAPID omics v2.0. Stacked bar plots (plots: **a**-**c**) show comparisons of iCLIP, HITS-CLIP and PAR-CLIP motif-containing regions (in percentage) against *CLIPper*_*Rep*._ (left panels) and *DEWSeq* (right panels) results. **Blue bars** depict percentage motifs containing regions exclusive to *CLIPper*_*Rep*._ results, **orange bars** depict percentage motifs containing regions exclusive to *DEWSeq* results and **gray bars** depict percentage of common motifs. (**a**) comparison of POSTAR2 iCLIP binding regions from Piranha and CIMS pipelines with eCLIP *CLIPper*_*Rep*._ results and *DEWSeq* results. (**b**) comparison of POSTAR2 HITS-CLIP binding regions from Piranha and CIMS pipelines with *CLIPper*_*Rep*._ results and *DEWSeq* results. (**c**) comparison of PAR-CLIP binding regions from Piranha, PARalyzer and Mukherjee^32^ pipelines with eCLIP *CLIPper*_*Rep*._ results and *DEWSeq* results. (**d**) Violin plots showing the absolute number of motif-containing regions in common either between iCLIP, HITS-CLIP, PAR-CLIP datasets analysed with *CIMS, Piranha, PARalyzer* or *Mukherjee* and ENCODE eCLIP data analysed with *CLIPper*_*Rep*._ and *DEWSeq*, respectively.

## Discussion

CLIP-mapped crosslink sites for RBPs frequently fall outside of their biological binding motifs. The crosslink sites of individual-nucleotide resolution CLIP methods such as eCLIP show significant variability of site distributions and locations relative to known sequence motifs bound by RNA-binding proteins (Fig. 2 and Supplementary Fig. 1). Notably, RBPs display accumulation of crosslink sites either centred on the motif (e.g. RBFOX2, hnRNPC), displaced to one side (e.g. CSTF2), immediately upstream (e.g. HNRNPA1, TROVE2), or immediately downstream (e.g. U2AF2) of the motif. Some also show crosslink site enrichment surrounding the motif, but depletion directly at the associated RNA sequence motif (e.g. HNRNPL, CPEB4) (Fig. 1d-f, Fig. 2 and Supplementary Fig. 1a-b). In the case of SLBP, which binds to 3’ UTR histone stem-loops^36^, crosslink sites accumulate upstream of the binding motif locations (Fig. 1c). However, the majority of CLIP protocols were primarily benchmarked on selected RBPs like RBFOX2 or hnRNPC which do show peak-like behaviour on top of the known target sites, justifying the choice of peak-callers for data analysis.

To increase robustness to the observed crosslinking patterns and therefore to improve the reliability of binding site identification, we developed a computational method that detects enriched regions of crosslink sites in single-nucleotide resolution CLIP called *DEWSeq*. Similar to *csaw* for ChIP-seq^27^, *DEWSeq* takes into account biological variation between replicates for significance testing of the IP samples against size-matched input (SMI) controls. We reanalysed 223 ENCODE eCLIP data sets covering 150 RBPs in one or both of two cell lines (K562 and HepG2), of which 107 RBPs had known experimentally determined RNA sequence motifs and one, SLBP, is known to target a specific RNA stem-loop secondary structure^17^. Using these sequence and structural motifs, we have performed, to the best of our knowledge, the most comprehensive eCLIP benchmarking study to date.

We showed that *DEWSeq*, even when operating on the minimal working requirement of two IP samples and one control sample, outperforms the single-replicate peak calling strategy of *CLIPper*_*Orig*._^*10*^ and *CLIPper*_*Rep*._^*14*^ (Fig. 3b). This is the case both for the number of motif-containing binding sites detected (a median 1.8-fold or 2.3 improvement) and for the percentage of sites that contain a motif, which approximates the true positive rate, for the majority of RBPs in the ENCODE data set (a median 7.1% or 3.9% improvement). *DEWSeq* discovers numerous motif-containing binding sites not found by *CLIPper*_*Rep*._, whereas CLIPper_*Rep*._ outperforms *DEWSeq* only in a handful of cases (Fig. 3d,e). Overall, *DEWSeq* increased the number of reported RBP-gene interactions 1.55-fold (median across RBPs and cell types) (Fig. 3c).

We used the RFAM covariance model for SLBP’s known histone 3’ UTR stem-loop structure targets^36^ to estimate accuracy and sensitivity of the binding site assignments. 39.2% of *DEWSeq*-identified binding regions contain the histone stem-loop, compared to 17.9% for *CLIPper*_*Rep*._. Interestingly, when searching the surrounding areas (up to 300 nt) of the crosslink sites, *DEWSeq* was able to detect >80% of all known stem-loops, whereas *CLIPper*_*Rep*._ levels off at ∼30%. *CLIPper*_*Rep*._ shows only minimal improvement even with 300 nt extensions to both sides. The far better exclusion of false-positives and an improvement in detecting true-positives suggests the superior potential of *DEWSeq* in identifying the target mRNA genes using secondary structure signals (Fig. 4c,d). Our benchmark of *DEWSeq* parameters highlights that bigger window sizes are beneficial (Supplementary Fig. 5a,b), however bigger window-sizes are not the driving factor for identifying true-positives in the case of SLBP (Fig. 4c,d). For SLBP, true positive rates are comparable between the methods and that extending the window around stem-loop structures does not lead to the detection of more binding sites for *CLIPper*_*Rep*._.

Although *DEWSeq* does have a minimal binding site length due to its window size parameter, for motif calling, *CLIPper* extends its binding sites by 50 base pairs upstream^14,41^. The reasoning is that the 5’ end of the *CLIPper* peak represents UV crosslink sites between protein and RNA, implying that the actual binding motif can be upstream. Crosslink site distributions around known RNA sequence motifs and secondary structures show differences in relative positions, up- or downstream of the target site, which justifies a broader searching frame.

*DEWSeq* does consistently improve the overlap with binding sites from other CLIP protocols compared to *CLIPper*_*Rep*._, although the overlap across protocols is very low overall (Fig. 5a-c). Our study includes a meta-analysis of binding sites generated using different protocols, methods and analysis tools. The general trend indicates that compared to iCLIP, HITS-CLIP and PAR-CLIP datasets, ENCODE eCLIP dataset analysed with *DEWSeq* contains a higher number of motif-containing binding sites. Also, binding sites discovered exclusively by *DEWSeq* can be found in other CLIP protocols, providing independent validation. However, a further large-scale investigation is needed to study the differences between CLIP-type protocols.

Where, as in ChIP-seq, bell-shape patterns are displayed in eCLIP or iCLIP crosslinking data, peak-callers should perform well. However, given the evidence provided by our analysis (Fig. 1b-f, Fig. 2 and Supplementary Fig. 1), eCLIP data can also show a general enrichment of crosslink sites adjacent to sequence motifs. Given the evidence we found for specific crosslink site patterns for well-known RBPs, we concluded that contrary to ChIP-seq, testing for enrichments of crosslink sites (IP over SMI control) with broader sliding windows is more appropriate for the analysis of eCLIP data. Though the underlying biology of the RNA-binding behaviour of the protein under investigation is key for understanding CLIP data, the reduction of false-positives and greatly increased number of motif-containing binding sites provided by *DEWSeq* should help improve functional analyses downstream. To fully capitalise on the potential of *DEWSeq*, we further highly recommend that any CLIP-type experiments should be performed with at least 3 replicates both for samples and controls, as this will drastically improve the statistical power for reliable binding site detection with *DEWSeq* in a way that standard eCLIP data processing using *CLIPper* cannot provide. *DEWSeq* is highly scalable, easy-to-use, open-source, fully documented and is designed to circumvent the limitations of the individual-nucleotide resolution CLIP protocols outlined above. Finally, based on the results shown, we strongly advise that CLIP-type protocols and analysis methods should be evaluated on RNA-binding proteins with a variety of crosslinking and binding behaviours, thereby taking into account structural and functional biological differences.

## Online Methods

### eCLIP data

In order to compare the results from our newly developed *DEWSeq* package, we chose the eCLIP data published by the ENCODE Project^10^. This dataset provides consistently produced 223 experimental studies, covering 150 RBPs in either one or both of the two human cell lines: HepG2 and K562. Each study in this dataset consists of two biological replicate IP samples and one size-matched input (SMI) control sample^42^. For data reanalysis using *DEWSeq*, we downloaded alignment files (.bam) with reads mapped to the GRCh38 genome annotations from ENCODE project data portal^43^. Bam file accessions and additional details are given in Supplementary Table 5.

### *CLIPper* binding sites

In this manuscript, the results from the reanalysis of this ENCODE dataset were compared against two sets of binding site results: the original set of called on individual IP samples with respect to SMI controls^10^ (referred to as *‘CLIPper original’ (CLIPper*_*Orig*._) in this manuscript) and the refined set of binding sites based on stringent thresholds, IDR analysis and blacklist region removal^14^ (referred to as ‘*CLIPper reproducible’* (*CLIPper*_*Rep*._) in this manuscript). Files providing these *CLIPper* results were also downloaded from the ENCODE Project data portal in narrowPeak BED format.

### Preprocessing of eCLIP data with *htseq-clip*

We have developed a custom python package called *htseq-clip*^*25*^ to count and extract crosslink sites from sequencing alignment files. *htseq-clip* is designed to preprocess alignment files from CLIP experiments and to generate a count matrix of crosslink sites that can be used in further downstream analysis. The required inputs for *htseq-clip* are: gene annotation in GFF format and alignment files (.bam format, coordinate sorted and indexed).

*htseq-clip* flattens the input gene annotation file and creates sliding windows by splitting each individual gene annotation feature into a series of overlapping (sliding) windows, where length and overlap (slide) are user supplied parameters. In subsequent steps, *htseq-clip* filters and extracts the crosslink sites based on user supplied experiment specifications and computes the number of crosslink sites per sliding window. In the final step, crosslink counts for multiple samples from the same experiment are summarised into a crosslink site count matrix file, which will be used as input for further downstream analysis. In this study, we used windows of size 50, 75 and 100 base pairs and slides of size 5 and 20 base pairs. The IP and SMI samples in each study were processed and concatenated into a crosslink site count matrix and was used as input for downstream analysis using *DEWSeq*.

### Calling differentially enriched regions for eCLIP data with *DEWSeq*

We developed the R/Bioconductor package *DEWSeq* to analyse the crosslink site count matrix from *htseq-clip* and to test for differential enrichment in IP samples in comparison with SMI samples. For statistical testing, *DEWSeq* utilises *DESeq2*^*26*^, a well-established R/Bioconductor package primarily used for the analysis of differentially expressed genes in RNA-seq data. After *DESeq2* initial pruning, normalisation and dispersion estimation, *DEWSeq* uses a custom one-tailed test for detecting significant crosslink regions enriched in IP samples over SMI, followed by two multiple hypothesis correction steps. In the first step, dependencies between overlapping windows are corrected using Bonferroni correction, as the adjacent sliding windows share crosslink site count information. In the second step, all windows are corrected for False Discovery Rates (FDR) at the genome level using either Benjamini-Hochberg (BH) method or independent hypothesis weighting (IHW)^44^. Finally, all enriched windows passing user specified enrichment thresholds are merged into regions.

*DEWSeq* is available as an open-source R/Bioconductor package (https://bioconductor.org/packages/release/bioc/vignettes/DEWSeq/inst/doc/DEWSeq.html). A sample pipeline for the analysis of eCLIP/iCLIP data using *htseq-clip* and *DEWSeq* is available^45^.

In this study, test for the impact of *DESeq2* and *DEWSeq* parameters on the final results, we ran *DEWSeq* with the following set of parameters on all ENCODE studies:

1. Dispersion estimation: Using *DESeq2* default “parametric” dispersion estimation^26^ or a custom function to decide the best fit (either “parametric” or “local” from *DESeq2*). Referred to either as “parametric” or “auto” throughout the rest of this manuscript.
2. Choice of statistical test: Either Wald test or Likelihood Ratio Test (LRT) from *DESeq2*. Referred to either as “Wald” or “LRT” throughout the rest of this manuscript.
3. Influence of dependent windows: either correct for dependencies between overlapping windows using Bonferroni correction or using no dependency correction. Referred to either as “Bonferroni” or “no correction” throughout the rest of this manuscript.
4. Choice of FDR correction methods: Either using Benjamini-Hochberg (BH) method for FDR correction or using IHW for FDR correction. Referred to either as “BH” or “IHW” throughout the rest of this manuscript.

All the enriched regions resulting from these parameter combinations were filtered using the following thresholds: log_2_ fold change ≥ 0.5 and p_adj_ value ≤ 0.1. Parameter combinations were benchmarked for robustness (Supplementary Fig. 4) and parameters 100 nt window size, 5 nt step size, LRT testing, “no correction”, IHW, and “auto fit” were selected. For the rest of this manuscript, these results will be referred to as *“DEWSeq binding sites”*.

### Gene annotations

We used all primary assembly gene models from GENCODE release 27 (GRCh38.p10) to map reported RBP binding sites to genes.

### Reference set of known RBP motifs

In this analysis, we used the presence of an RNA sequence motif in a binding region as a proxy for the biological relevance of the binding region. We used motifs from the catRAPID omics v2.0 RBP motifs database, whose authors collected and curated motifs from a comprehensive set of sources including ATtRACT, cisBP-RNA, mCrossBase, oRNAment and RBPmap^29^. Supplementary Table. 5, Sheet 2 shows the total number of RBPs per motif length in the data source and the number of RBPs in common with the ENCODE dataset.

We post-processed this set of motifs by removing any peripheral positions with information content ≤ 0.1 and rounding down base probabilities ≤ 0.025 using the R/Bioconductor package universalmotif^46^. We then selected motifs with length ≥ 6 nt to reduce the probability of random occurrences. This selected set of motifs comprised 604 motifs from 258 RBPs, 107 of which had ENCODE eCLIP data available. Using motifs from this common set of RBPs, our final benchmark set contained 322 motifs for 107 RBPs (Supplementary File 1).

### Discovery and comparison of known motifs sites

We predicted the positions of human RNA-binding protein motifs within eCLIP binding regions using version 5.4.0 of FIMO^30^ from the MEME Suite^47^ and the comprehensive benchmark set of RBP motifs described above. For CLIPper, eCLIP binding regions were extended 50 nt upstream of their 5’ end, as previously described^14,41^. Enriched regions from *DEWSeq* were analysed without any extension. We used a near-equiprobable background sequence model calculated across GENCODE 27 transcripts using fasta-get-markov (-norc) from the MEME Suite. We filtered FIMO results using a p-value cutoff ≤ 0.001. We consciously avoided the use of q-values to filter the results due to the variation in the number of binding sites between different RBPs, which would penalise experiments that succeeded at identifying a larger number of binding sites, and reasoned that p-values would offer more comparable results across experiments.

### Secondary structure analysis

This analysis was restricted to enriched regions from SLBP, as it was the only RBP in the ENCODE data with well characterised RNA secondary structure binding targets (histone 3’ UTR stem-loops) to the best of our knowledge. In this study, we used *cmsearch* tool from Infernal suite^48^ and covariance models from Rfam database^49^ to scan for secondary structures in enriched regions from both *CLIPper*_*Rep*._ and *DEWSeq* results. In the first step, we used *mergeBed* from bedtools suite^50^ to merge overlapping regions or regions separated by a maximum of 10 nucleotides in *CLIPper*_*Rep*._ SLBP results. In case of *DEWSeq* SLBP regions, no such merging step was necessary. In the next step, we sequentially extended the length of both *DEWSeq* and *CLIPper*_*Rep*._ regions by up to 300 nt (50, 100, 150, 200, 300) using *slopBed* tool from the bedtool suite. With this extension step, we aim to detect histone stem-loop structures that are found in close proximity to either *CLIPper*_*Rep*._ or *DEWSeq* enriched regions, and to assess the gain in number of stem-loop structures identified with each extension. The fasta sequences extracted (using *getfasta* from bedtools) from the original set of regions and extended regions were subjected to a profile-based search with the histone 3′ UTR stem-loop family (Rfam id: RF00032) covariance model and using Infernal *cmsearch* tool. In this step, *cmsearch* sequence-based pre-filtering heuristics were turned off and an E-value threshold ≥ 5.0 was used to identify hits. The *cmsearch* output table was processed with bash *awk* command to obtain the genomic location of the model hits and extracted unique hits based on the genomic coordinates. The reference set of Histone 3’ UTR stem-loop regions were retrieved from the Rfam database (Rfam id: RF00032).

### Comparison to orthogonal datasets

We retrieved iCLIP^15^, HITS-CLIP^7^, and PAR-CLIP^8^ interaction datasets from the POSTAR2 database (previously known as CLIPdb)^51^. For each CLIP experiment type, POSTAR2 provided peak calling results from *Piranha*^*38*^, as well as the more specialised *CIMS* (crosslink-induced mutation sites)^39^ and *PARalyzer* data analysis pipelines^40^. We used POSTAR2’s standard thresholds such as: p-value < 0.01 for *Piranha*, score < 0.01 for *CIMS*, and score > 0.5 for *PARalyzer*. Additionally, we also obtained a cohesive PAR-CLIP dataset from the DoRiNA database ^32^, and used a minimum conversion specificity score of 5 to filter binding regions, which resulted in a similar number of regions as for *CLIPper*_*Rep*._. We performed motif calling on these enriched regions using the reference motif set and methodology described above and compared the fraction of unique motifs present in these results to that of *DEWSeq* and *CLIPper*_*Rep*._ results.

### Comparing gene region and gene type enrichment using OLOGRAM

To determine whether *DEWSeq* or *CLIPper* results were biassed towards gene regions (5’ UTR, exon, 3’ UTR) or gene types (e.g. protein coding RNAs, non coding RNA, mtRNA, …) we performed gene region and gene type enrichment analysis in both results using OLOGRAM^52^. To avoid ambiguities in gene region annotation in genes with multiple transcripts, we selected the transcript with the highest abundance in each gene as the representative transcript. For this purpose we used rRNA-depleted total RNA-seq data from HepG2 (ENCODE accession: ENCFF533XPJ, ENCFF321JIT) and K562 (ENCODE accession: ENCFF286GLL, ENCFF986DBN) cell lines available from ENCODE consortium. After removing lowly expressed transcripts with a TPM value ≤ 1, the datasets were merged and for each gene, the transcript with the highest abundance was selected as the representative transcript. We selected 15,274 transcripts as candidates and extracted region and type annotation for these selected transcripts. In the next step, we used this gene annotation data and enriched regions from *DEWSeq* and *CLIPper*_*Rep*._ results to assess the significance of overlaps between enriched regions in either of the two sets and the gene region or gene type annotations.

## Supporting information

Supplementary File 1 - Motifs

Supplementary File 2 - DEWSeq Binding Sites

## Availability

*DEWSeq* is available as an R/Bioconductor package. Reported *DEWSeq* binding sites are available as Supplementary File 2.

## Acknowledgements

We thank Alessio Colantoni and Alexandros Armaos for providing the curated RBP motif dataset from Armaos *et al*. (2021). We thank Thileepan Sekaran for bioinformatics help.

## Supplementary Material

**Supplementary Table 1** | Results of RNA sequence motif benchmark on RNA-binding protein binding site detections methods

**Supplementary Table 2** | Results of motif exclusiveness analysis, comparison to iCLIP, HITS-CLIP and PAR-CLIP datasets (Sheet 1), Overlap of iCLIP, HITS-CLIP, PAR-CLIP datasets from POSTAR with ENCODE eCLIP (Sheet 2)

**Supplementary Table 3** | RBFOX2 UGCAUG motif counts (Sheet 1), Enrichment analysis results across all chromosomes for all FASTKD2 samples (Sheet 2)

**Supplementary Table 4** | Analysis of histone stem-loop found in SLBP eCLIP binding sites and extensions to 5’ and 3’ end.

**Supplementary Table 5** | Details of ENCODE eCLIP data used (Sheet 1). Total number of RBPs per motif length in the data source and the number of RBPs in common with the ENCODE dataset (Sheet 2).

**Supplementary File 1** | RBP motifs in .meme file format and as motif logos.

**Supplementary File 2** | DEWSeq enriched binding regions in narrowPeak format

**Supplementary Figure 1.**
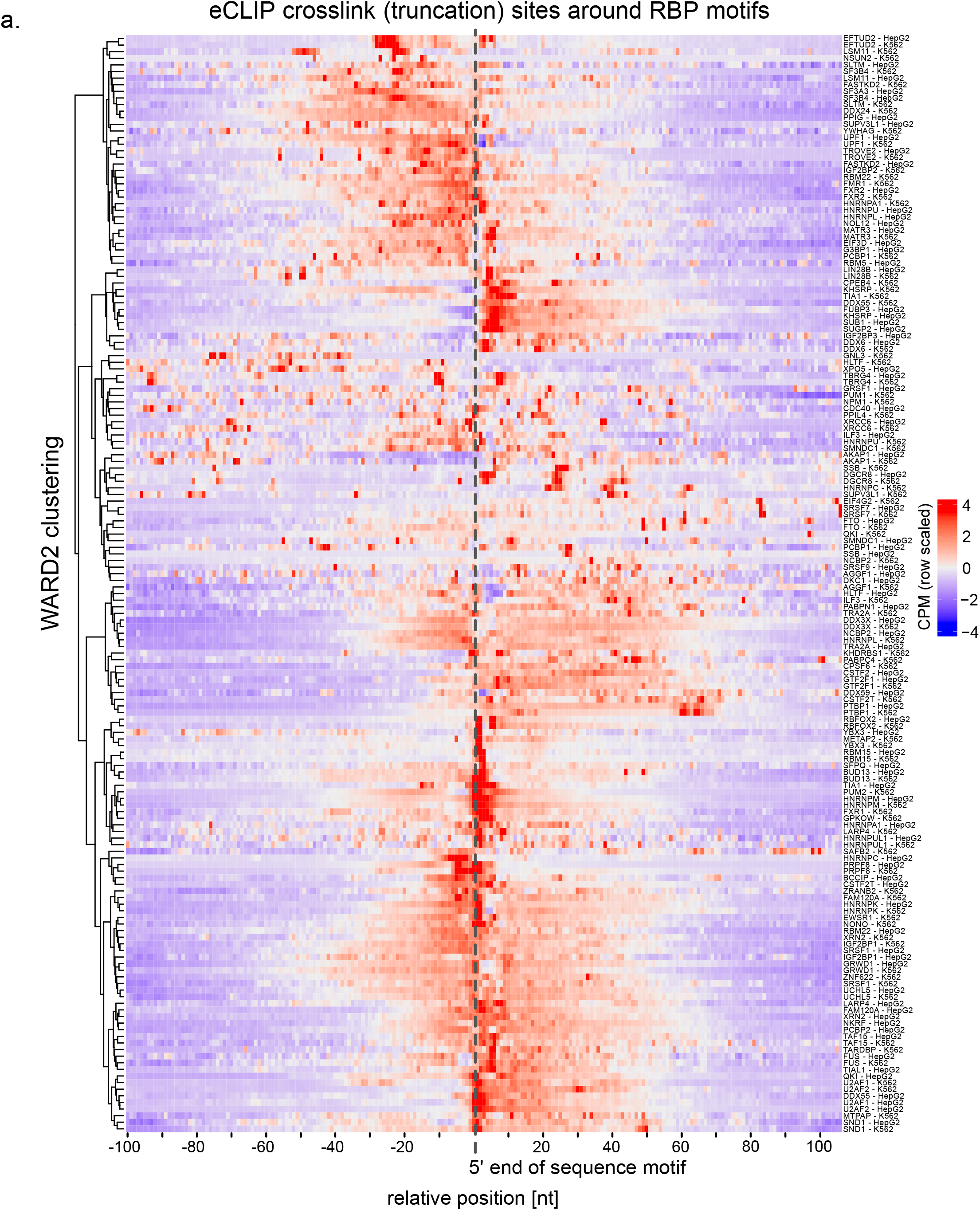
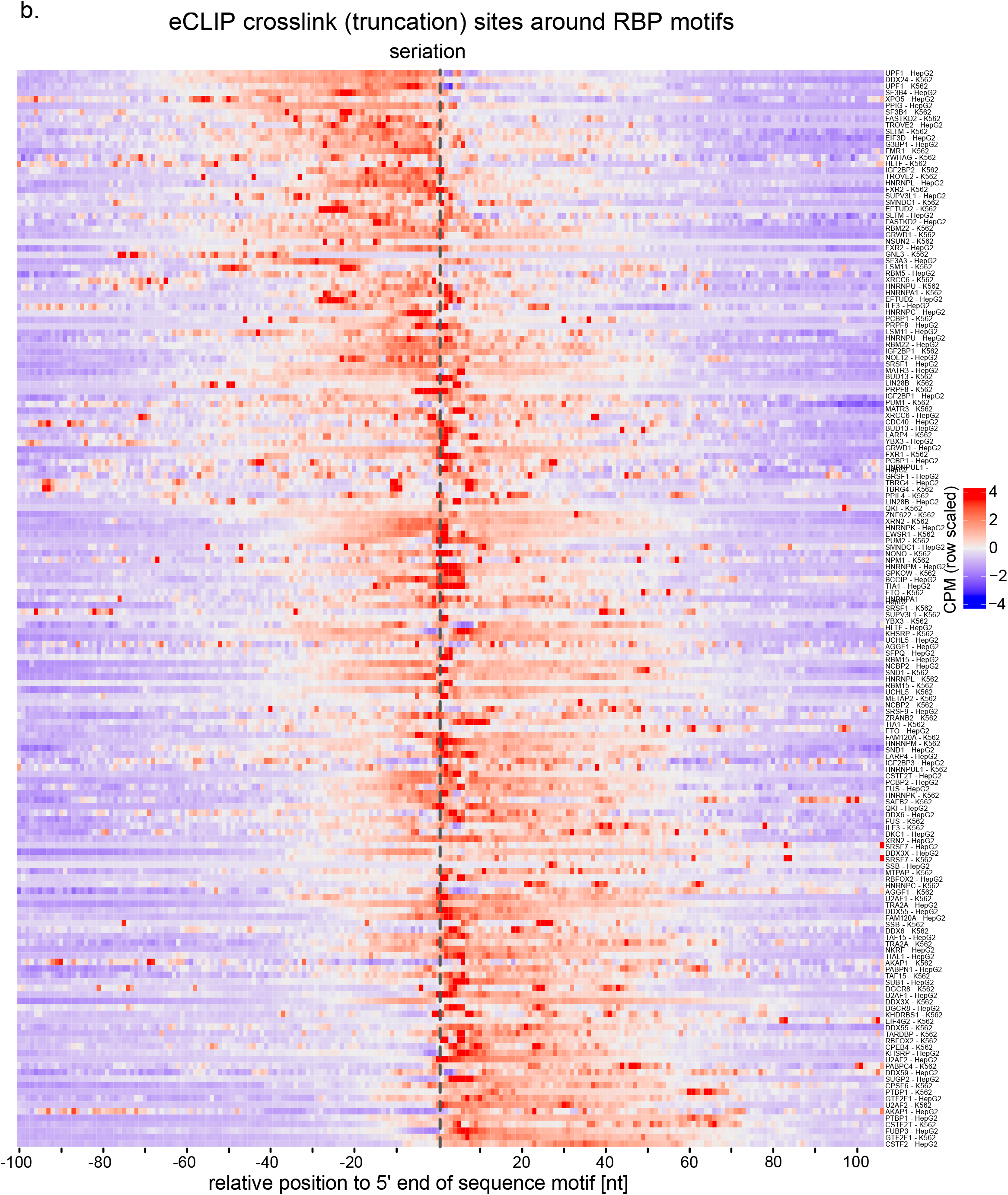
Heatmaps of crosslink sites around experimentally derived RNA sequence motifs. Each row displays a motif for an RNA-binding protein. (**a**) clustered with WARD2 to group similar binding patterns and (**b**) clustered with seriation to reveal the broader structure of crosslinking sites around motifs.

**Supplementary Figure 2.**
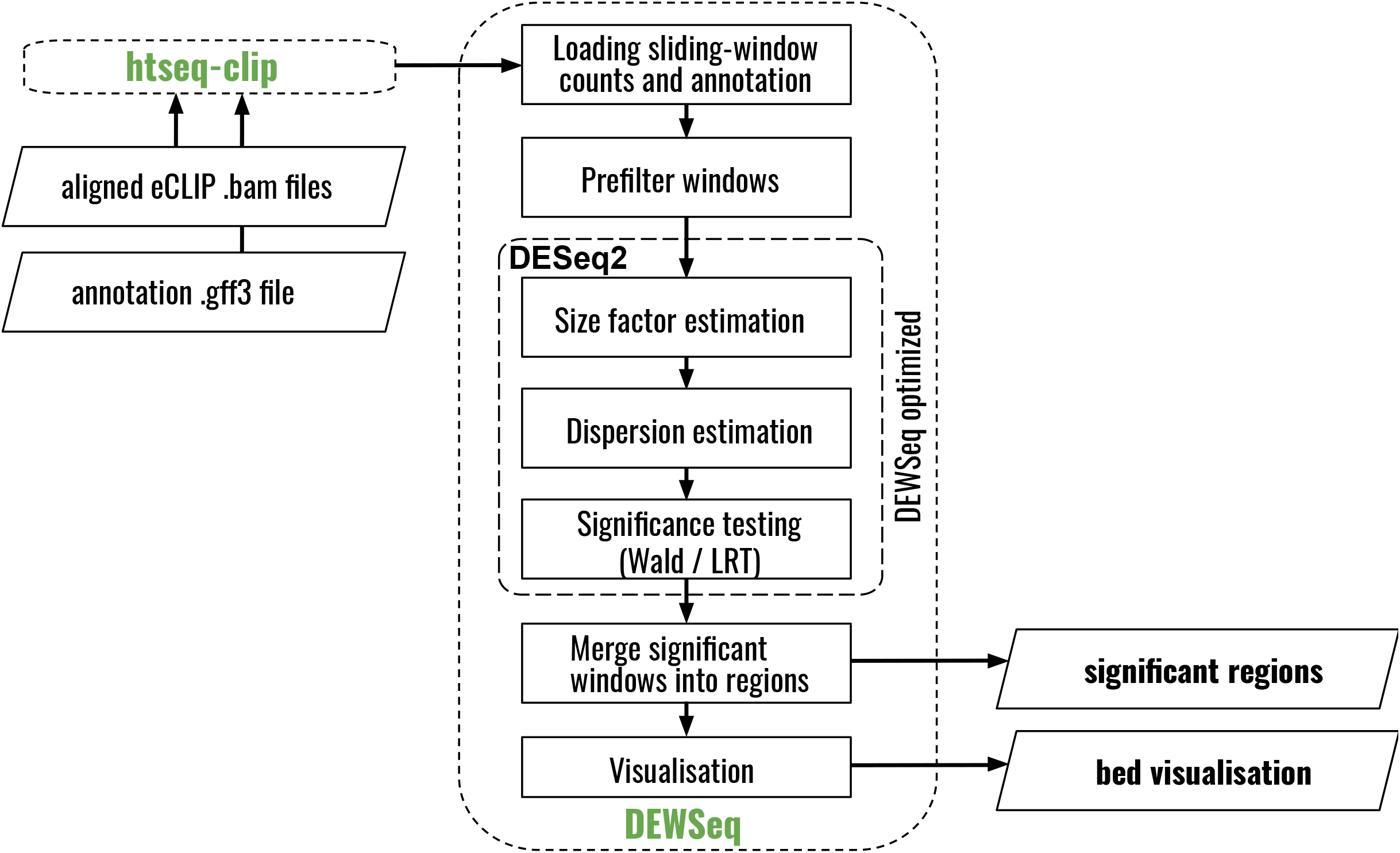
Overview of DEWSeq workflow

**Supplementary Figure 3.**
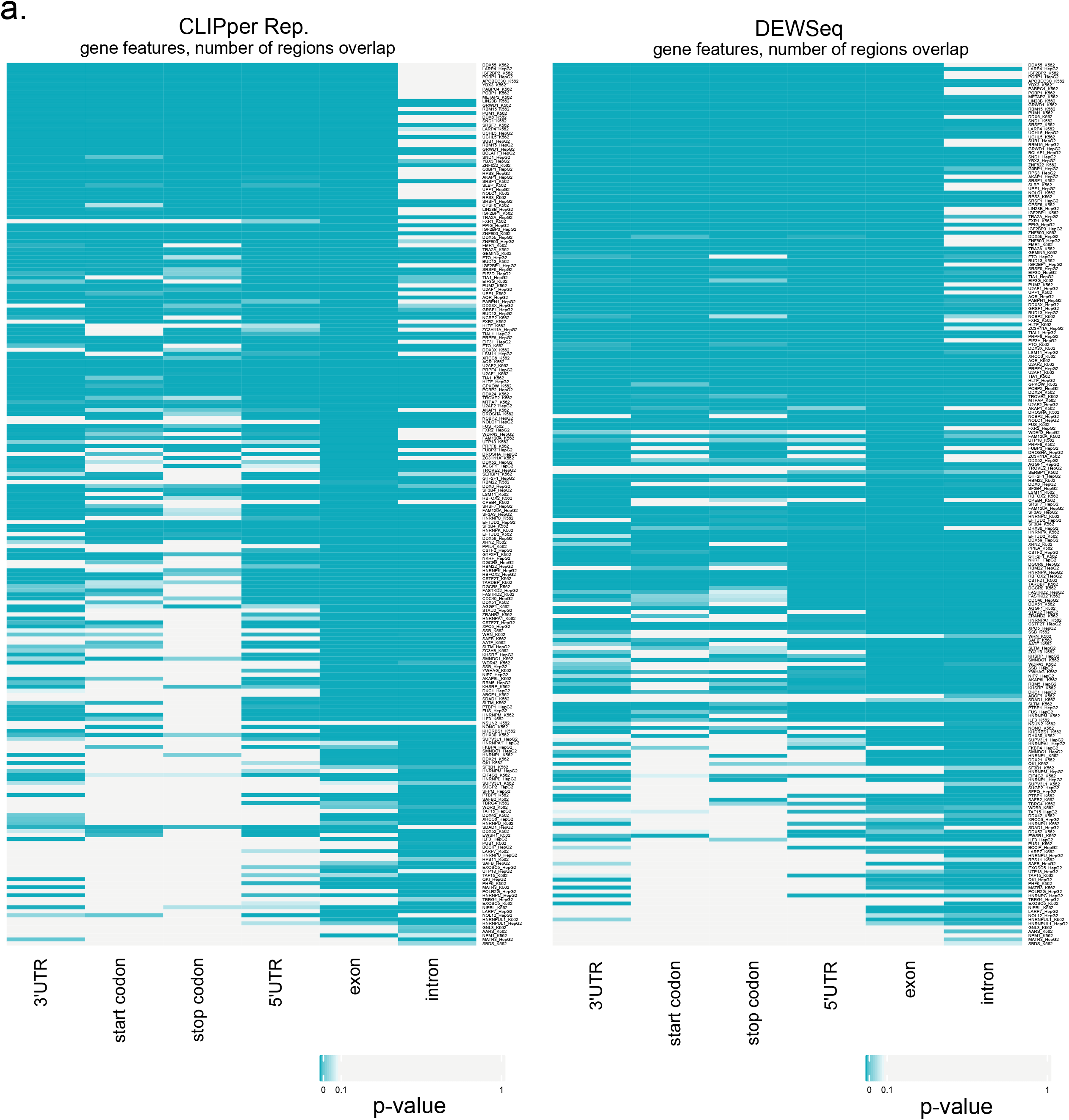
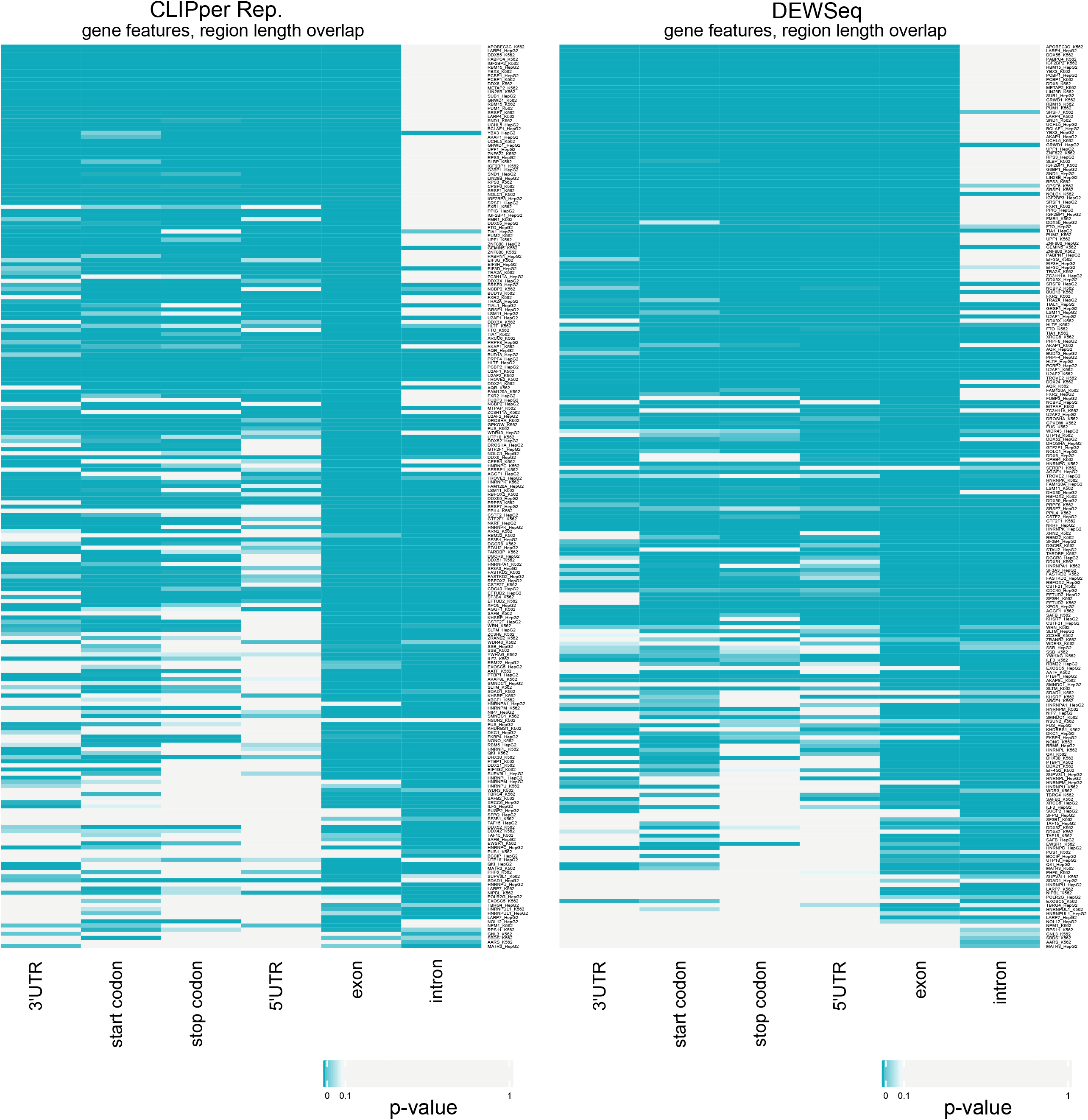
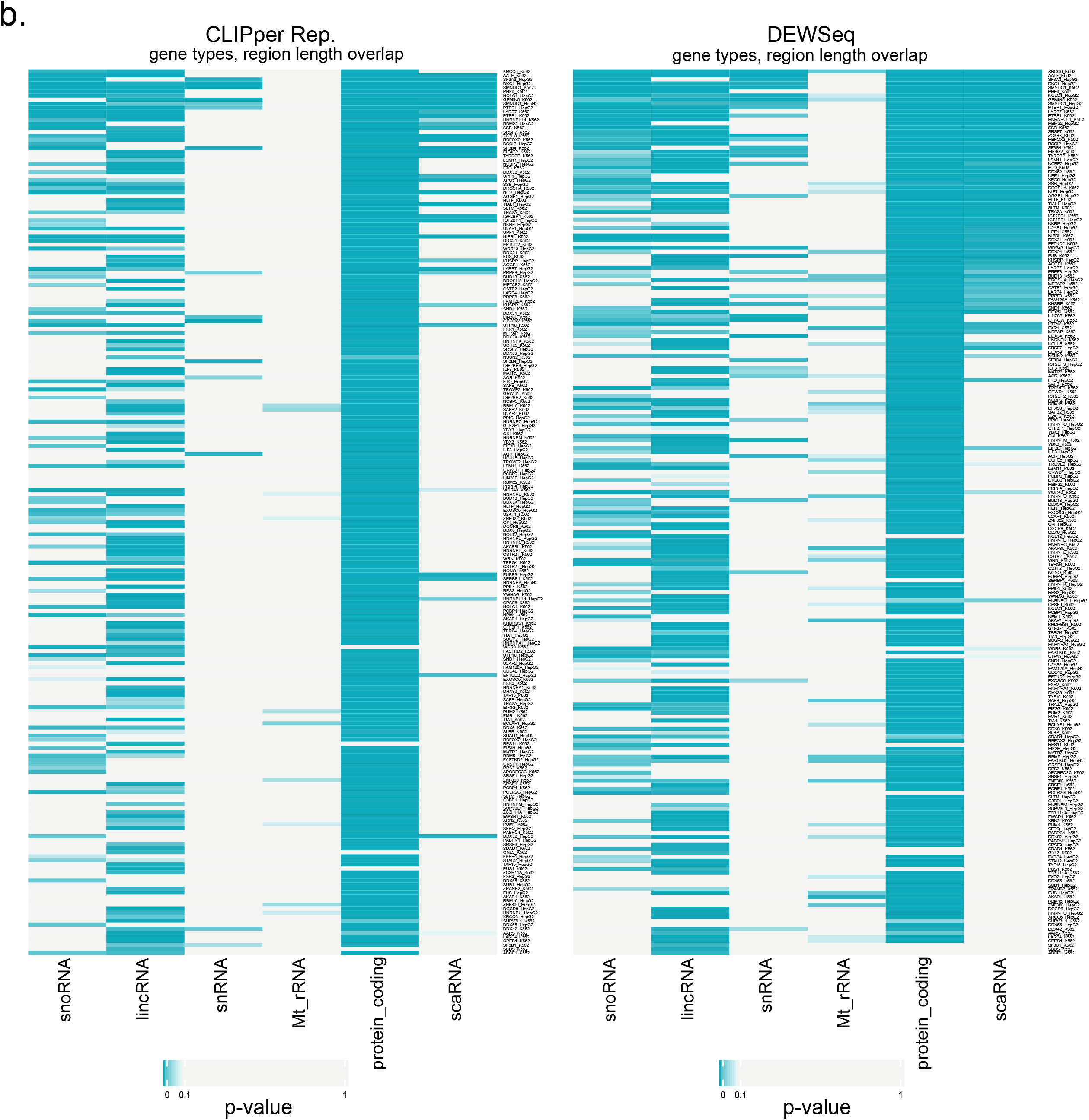
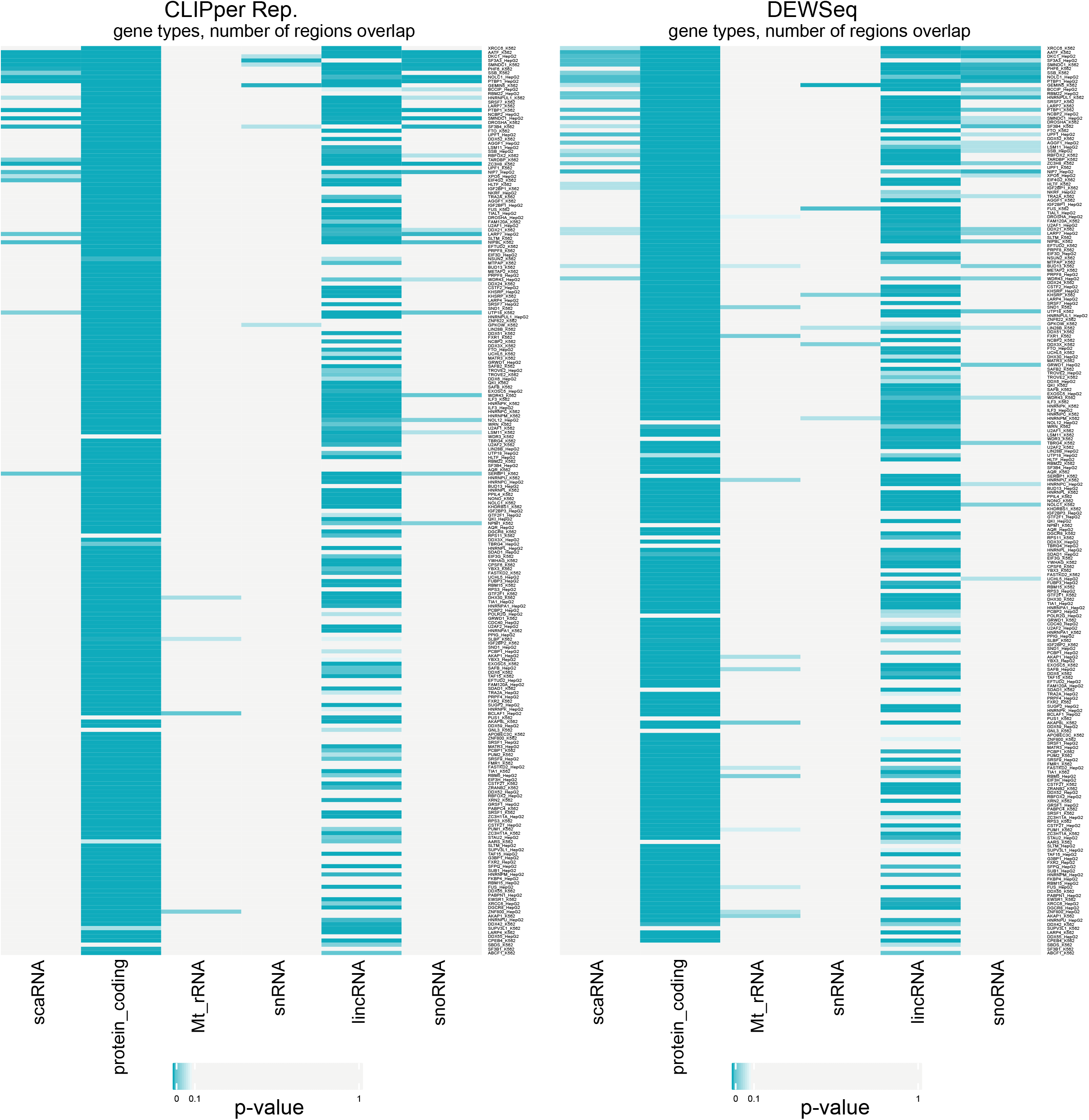
Functional analysis of (**a**) gene feature or (**b**) gene type enrichment of ‘CLIPper reproducible’ (*CLIPper*_*Rep*._) and DEWSeq binding sites with OLOGRAMS.

**Supplementary Figure 4.**
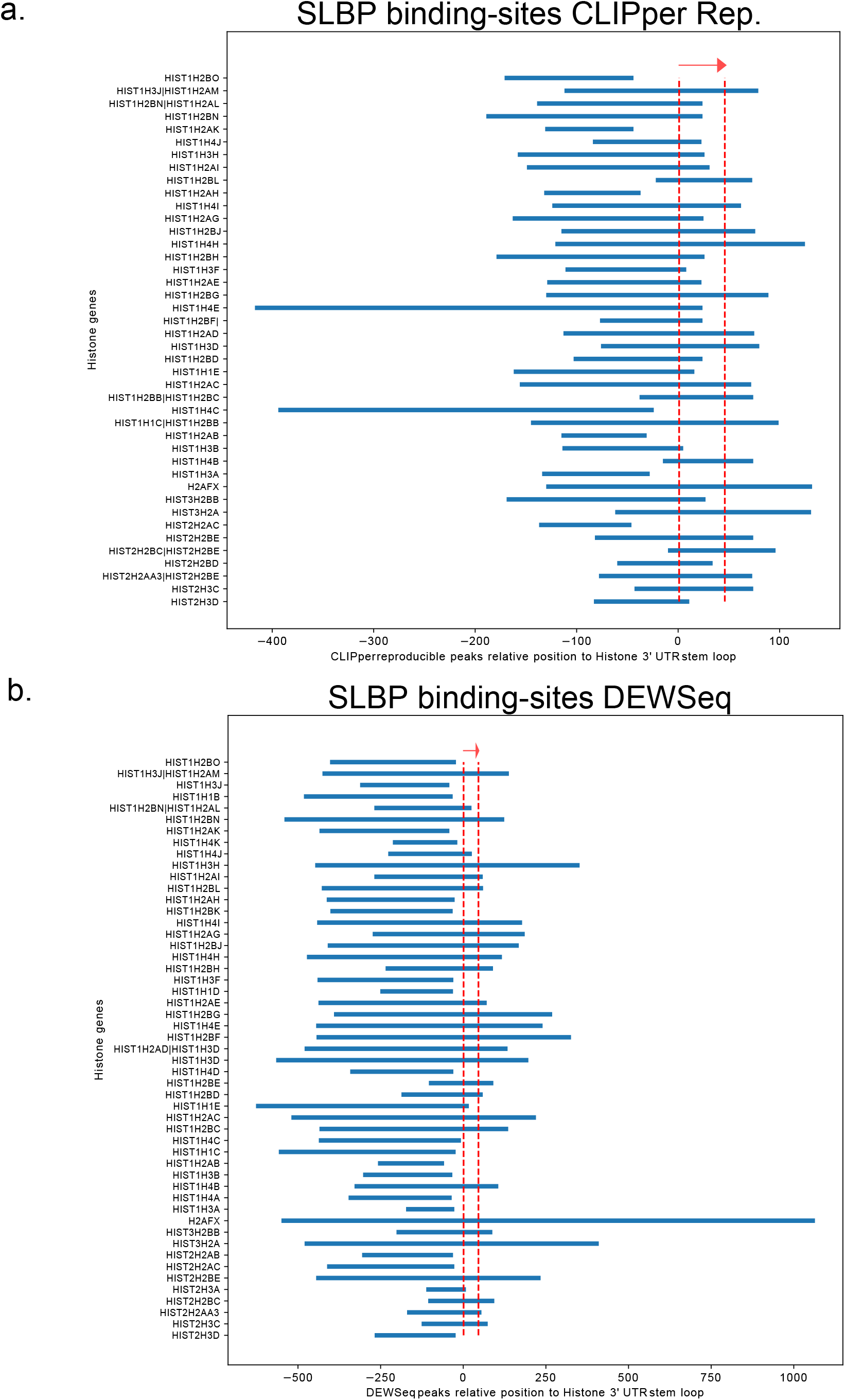
SLBP eCLIP binding sites detected by (**a**) ‘CLIPper reproducible’ and (**b**) DEWSeq positions respective to the known 3’ histone mRNA stem-loop targets

**Supplementary Figure 5.**
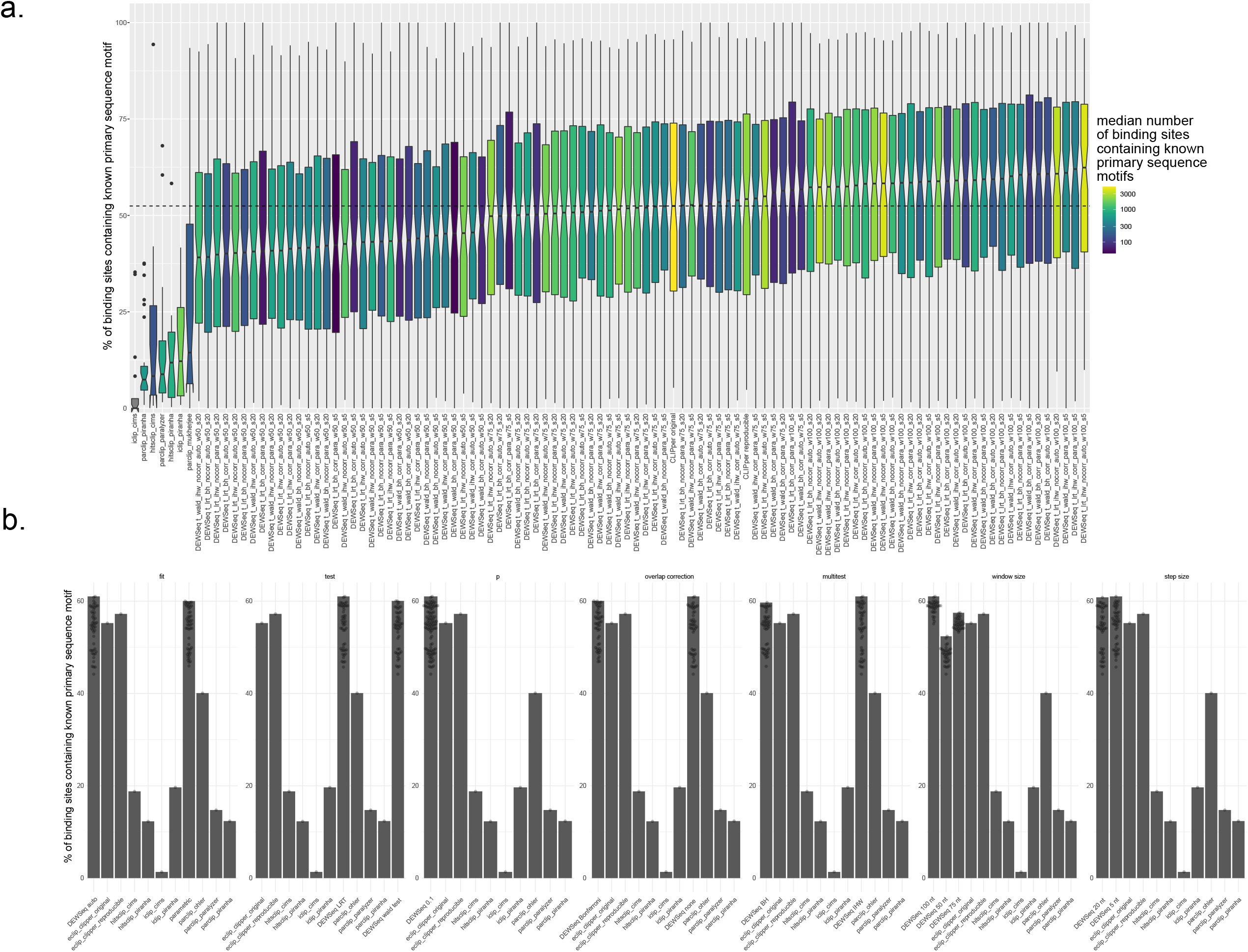
Test of different DEWSeq parameters (**a**) in combination (**b**) for robustness

**Supplementary Figure 6.**
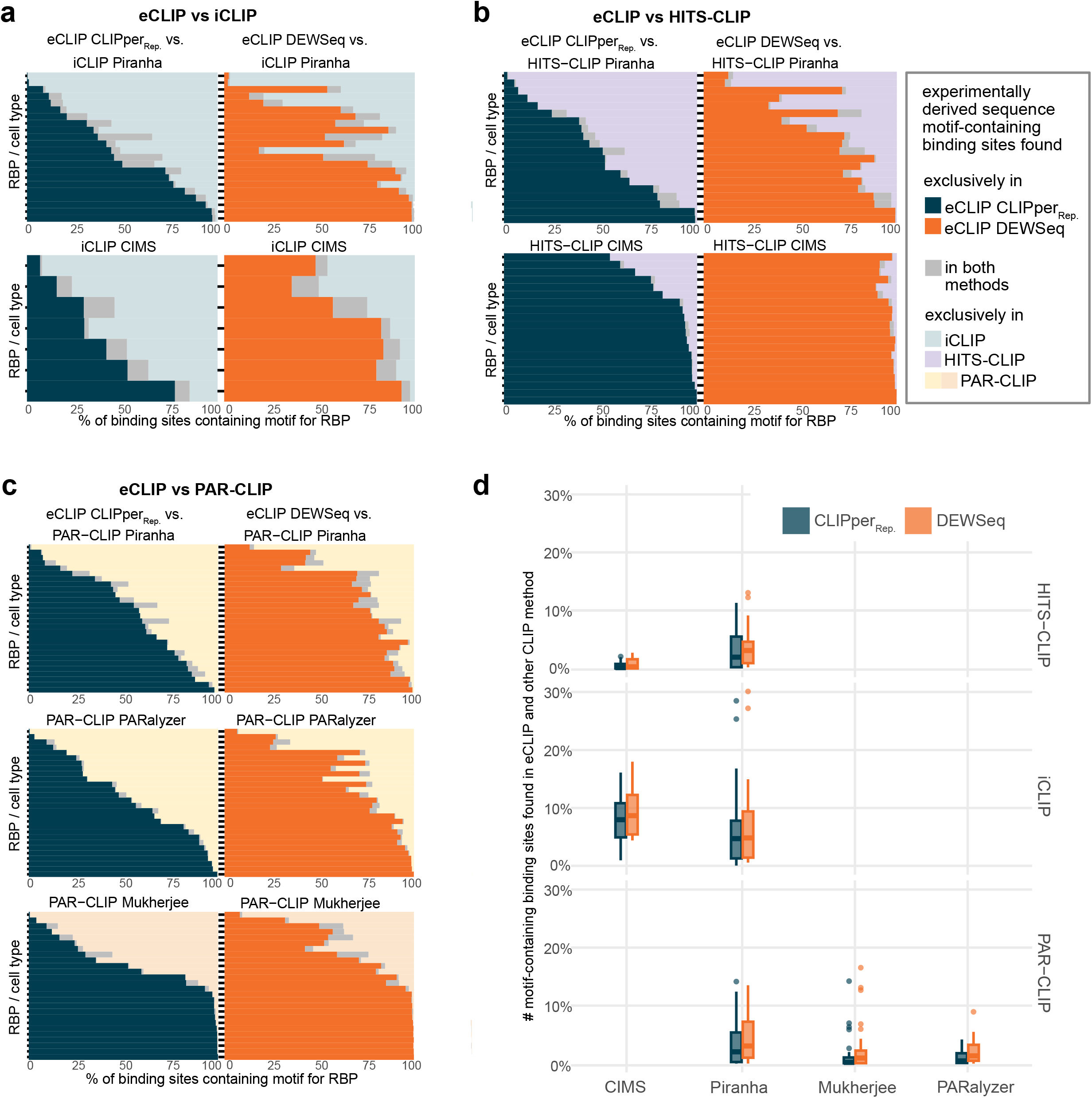
Binding site exclusiveness. Comparison of motif-containing binding sites from various CLIP datasets against eCLIP *‘CLIPper reproducible’* (*CLIPper*_*Rep*._) and *DEWSeq* results. The comparisons are for common RBPs (in ENCODE dataset and these methods) with motifs in catRAPID omics v2.0. Stacked bar plots (plots: **a**-**c**) show comparisons of iCLIP, HITS-CLIP and PAR-CLIP motif-containing regions (in percentage) against *CLIPper*_*Rep*._ (left panels) and *DEWSeq* (right panels) results. Blue bars depict percentage motifs containing regions exclusive to *CLIPper*_*Rep*._ results, orange bars depict percentage motifs containing regions exclusive to *DEWSeq* results and gray bars depict percentage of common motifs. RBPs for *CLIPper*_*Rep*._ and *DEWSeq* are in the same order. (**a**) comparison of POSTAR2 iCLIP binding regions from Piranha and CIMS pipelines with eCLIP *CLIPper*_*Rep*._ results and *DEWSeq* results. (**b**) comparison of POSTAR2 HITS-CLIP binding regions from Piranha and CIMS pipelines with *CLIPper*_*Rep*._ results and *DEWSeq* results. (**c**) comparison of PAR-CLIP binding regions from Piranha, PARalyzer and Mukherjee^32^ pipelines with eCLIP *CLIPper*_*Rep*._ results and *DEWSeq* results. (**d**) Boxplots showing the percentage of motif-containing regions in common either between iCLIP, HITS-CLIP, PAR-CLIP datasets analysed with *CIMS, Piranha, PARalyzer* or *Mukherjee* and ENCODE eCLIP data analysed with *CLIPper*_*Rep*._ and *DEWSeq*, respectively.

## Abbreviations

adj: adjusted
Orig: original
IP: immunoprecipitated
CIMS: crosslink-induced mutation sites
CLIP: crosslinking immunoprecipitation
nt: nucleotide(s)
PAR: photoactivatable ribonucleoside enhanced
Rep: reproducible
SMI: size-matched input
TPM: tags per million
UTR: untranslated region

## References

1. Ule, J. & Blencowe, B. J. Alternative Splicing Regulatory Networks: Functions, Mechanisms, and Evolution. Mol. Cell 76, 329–345 (2019).

2. Glisovic, T., Bachorik, J. L., Yong, J. & Dreyfuss, G. RNA-binding proteins and post-transcriptional gene regulation. FEBS Letters vol. 582 1977–1986 Preprint at https://doi.org/10.1016/j.febslet.2008.03.004 (2008).

3. Gebauer, F., Schwarzl, T., Valcárcel, J. & Hentze, M. W. RNA-binding proteins in human genetic disease. Nat. Rev. Genet. 22, 185–198 (2021).

4. Hentze, M. W., Castello, A., Schwarzl, T. & Preiss, T. A brave new world of RNA-binding proteins. Nat. Rev. Mol. Cell Biol. 19, 327–341 (2018).

5. Hafner, M. et al. CLIP and complementary methods. Nature Reviews Methods Primers 1, 1–23 (2021).

6. Ule, J. et al. CLIP identifies Nova-regulated RNA networks in the brain. Science 302, 1212–1215 (2003).

7. Licatalosi, D. D. et al. HITS-CLIP yields genome-wide insights into brain alternative RNA processing. Nature 456, 464–469 (2008).

8. Hafner, M. et al. Transcriptome-wide identification of RNA-binding protein and microRNA target sites by PAR-CLIP. Cell 141, 129–141 (2010).

9. König, J., Zarnack, K., Luscombe, N. M. & Ule, J. Protein–RNA interactions: new genomic technologies and perspectives. Nature Reviews Genetics vol. 13 77–83 Preprint at https://doi.org/10.1038/nrg3141 (2012).

10. Van Nostrand, E. L. et al. Robust transcriptome-wide discovery of RNA-binding protein binding sites with enhanced CLIP (eCLIP). Nat. Methods 13, 508–514 (2016).

11. Zarnegar, B. J. et al. irCLIP platform for efficient characterization of protein–RNA interactions. Nat. Methods 13, 489–492 (2016).

12. Van Nostrand, E. L. et al. Robust, Cost-Effective Profiling of RNA Binding Protein Targets with Single-end Enhanced Crosslinking and Immunoprecipitation (seCLIP). Methods Mol. Biol. 1648, 177–200 (2017).

13. Porter, D. F. et al. easyCLIP analysis of RNA-protein interactions incorporating absolute quantification. Nat. Commun. 12, 1569 (2021).

14. Van Nostrand, E. L. et al. A large-scale binding and functional map of human RNA-binding proteins. Nature 583, 711–719 (2020).

15. Konig, J. et al. iCLIP - Transcriptome-wide Mapping of Protein-RNA Interactions with Individual Nucleotide Resolution. Journal of Visualized Experiments Preprint at https://doi.org/10.3791/2638 (2011).

16. Zarnack, K. et al. Direct competition between hnRNP C and U2AF65 protects the transcriptome from the exonization of Alu elements. Cell 152, 453–466 (2013).

17. Dominski, Z., Zheng, L. X., Sanchez, R. & Marzluff, W. F. Stem-loop binding protein facilitates 3’-end formation by stabilizing U7 snRNP binding to histone pre-mRNA. Mol. Cell. Biol. 19, 3561–3570 (1999).

18. Nourse, J., Spada, S. & Danckwardt, S. Emerging Roles of RNA 3′-end Cleavage and Polyadenylation in Pathogenesis, Diagnosis and Therapy of Human Disorders. Biomolecules vol. 10 915 Preprint at https://doi.org/10.3390/biom10060915 (2020).

19. Mackereth, C. D. et al. Multi-domain conformational selection underlies pre-mRNA splicing regulation by U2AF. Nature 475, 408–411 (2011).

20. Smith, S. A. et al. Paralogs hnRNP L and hnRNP LL exhibit overlapping but distinct RNA binding constraints. PLoS One 8, e80701 (2013).

21. Schelhorn, C. et al. RNA recognition and self-association of CPEB4 is mediated by its tandem RRM domains. Nucleic Acids Res. 42, 10185–10195 (2014).

22. Huppertz, I. et al. RNA regulates Glycolysis and Embryonic Stem Cell Differentiation via Enolase 1. Cold Spring Harbor Laboratory 2020.10.14.337444 (2020) doi:10.1101/2020.10.14.337444.

23. Hauer, C. et al. Improved binding site assignment by high-resolution mapping of RNA-protein interactions using iCLIP. Nat. Commun. 6, 7921 (2015).

24. Hauer, C. et al. Exon Junction Complexes Show a Distributional Bias toward Alternatively Spliced mRNAs and against mRNAs Coding for Ribosomal Proteins. Cell Rep. 16, 1588–1603 (2016).

25. Sahadevan, S. et al. Htseq-clip: a toolset for the preprocessing of eCLIP/iCLIP datasets. Bioinformatics (2022) doi:10.1093/bioinformatics/btac747.

26. Love, M. I., Huber, W. & Anders, S. Moderated estimation of fold change and dispersion for RNA-seq data with DESeq2. Genome Biol. 15, 550 (2014).

27. Lun, A. T. L. & Smyth, G. K. csaw: a Bioconductor package for differential binding analysis of ChIP-seq data using sliding windows. Nucleic Acids Res. 44, e45 (2016).

28. Wheeler, E. C., Van Nostrand, E. L. & Yeo, G. W. Advances and challenges in the detection of transcriptome-wide protein-RNA interactions. Wiley Interdiscip. Rev. RNA 9, (2018).

29. Armaos, A., Colantoni, A., Proietti, G., Rupert, J. & Tartaglia, G. G. catRAPID omics v2.0: going deeper and wider in the prediction of protein–RNA interactions. Nucleic Acids Res. 49, W72–W79 (2021).

30. Grant, C. E., Bailey, T. L. & Noble, W. S. FIMO: scanning for occurrences of a given motif. Bioinformatics 27, 1017–1018 (2011).

31. Zhu, Y. et al. POSTAR2: deciphering the post-transcriptional regulatory logics. Nucleic Acids Res. 47, D203–D211 (2019).

32. Mukherjee, N. et al. Deciphering human ribonucleoprotein regulatory networks. Nucleic Acids Res. 47, 570–581 (2019).

33. Kuroyanagi, H. Fox-1 family of RNA-binding proteins. Cell. Mol. Life Sci. 66, 3895–3907 (2009).

34. Popow, J. et al. FASTKD2 is an RNA-binding protein required for mitochondrial RNA processing and translation. RNA 21, 1873–1884 (2015).

35. Jourdain, A. A. et al. A mitochondria-specific isoform of FASTK is present in mitochondrial RNA granules and regulates gene expression and function. Cell Rep. 10, 1110–1121 (2015).

36. Wang, Z. F., Whitfield, M. L., Ingledue, T. C., 3rd, Dominski, Z. & Marzluff, W. F. The protein that binds the 3’ end of histone mRNA: a novel RNA-binding protein required for histone pre-mRNA processing. Genes Dev. 10, 3028–3040 (1996).

37. Zanier, K. et al. Structure of the histone mRNA hairpin required for cell cycle regulation of histone gene expression. RNA 8, 29–46 (2002).

38. Uren, P. J. et al. Site identification in high-throughput RNA–protein interaction data. Bioinformatics vol. 28 3013–3020 Preprint at https://doi.org/10.1093/bioinformatics/bts569 (2012).

39. Moore, M. J. et al. Mapping Argonaute and conventional RNA-binding protein interactions with RNA at single-nucleotide resolution using HITS-CLIP and CIMS analysis. Nature Protocols vol. 9 263–293 Preprint at https://doi.org/10.1038/nprot.2014.012 (2014).

40. Corcoran, D. L. et al. PARalyzer: definition of RNA binding sites from PAR-CLIP short-read sequence data. Genome Biol. 12, R79 (2011).

41. Dominguez, D. et al. Sequence, Structure, and Context Preferences of Human RNA Binding Proteins. Mol. Cell 70, 854–867.e9 (2018).

42. Van Nostrand, E. L. et al. Principles of RNA processing from analysis of enhanced CLIP maps for 150 RNA binding proteins. Genome Biol. 21, 90 (2020).

43. Sloan, C. A. et al. ENCODE data at the ENCODE portal. Nucleic Acids Research vol. 44 D726–D732 Preprint at https://doi.org/10.1093/nar/gkv1160 (2016).

44. Ignatiadis, N., Klaus, B., Zaugg, J. B. & Huber, W. Data-driven hypothesis weighting increases detection power in genome-scale multiple testing. Nat. Methods 13, 577–580 (2016).

45. Sahadevan, S., Sekaran, T. & Schwarzl, T. A Pipeline for Analyzing eCLIP and iCLIP Data with Htseq-clip and DEWSeq. Methods Mol. Biol. 2404, 189–205 (2022).

46. Tremblay, B. J. M. universalmotif: Import. Modify, and Export Motifs with R (2020).

47. Bailey, T. L. et al. MEME SUITE: tools for motif discovery and searching. Nucleic Acids Res. 37, W202–8 (2009).

48. Nawrocki, E. P., Kolbe, D. L. & Eddy, S. R. Infernal 1.0: inference of RNA alignments. Bioinformatics 25, 1335–1337 (2009).

49. Kalvari, I. et al. Rfam 14: expanded coverage of metagenomic, viral and microRNA families. Nucleic Acids Res. 49, D192–D200 (2021).

50. Quinlan, A. R. & Hall, I. M. BEDTools: a flexible suite of utilities for comparing genomic features. Bioinformatics 26, 841–842 (2010).

51. Hu, B., Yang, Y.-C. T., Huang, Y., Zhu, Y. & Lu, Z. J. POSTAR: a platform for exploring post-transcriptional regulation coordinated by RNA-binding proteins. Nucleic Acids Res. 45, D104–D114 (2017).

52. Ferré, Q. et al. OLOGRAM: Determining significance of total overlap length between genomic regions sets. Bioinformatics (2019) doi:10.1093/bioinformatics/btz810.

